# Mild and Reversible Proprioception Perturbation Suggests Causal Biomechanics for Memory-Dependent Spatial Behavior in Mice

**DOI:** 10.64898/2026.07.28.740838

**Authors:** Meng-Xuan Liu, Nikky Chia-Ni Chang, Abe Ernest Johann Estalilla Isagan, Cheng-Han Lee, Ming-Yuan Min, Chih-Cheng Chen, Ching-Lung Hsu

**Author notes:** **Lead Contact:** Ching-Lung Hsu.

## Abstract

The neural system at the periphery is a frontline for embodied cognition, yet an acute, mild perturbation to dissect functional causality is hard to achieve. Peripheral neural processes and the central nervous system may work in concert to generate sophisticated representations regarding self and environments in the brain. This hypothesis, together with the underlying mechanisms, is particularly difficult to test for certain sensory inputs due to the lack of reversible manipulation techniques. Long postulated as a component for path integration, proprioception is one of such modalities. In this study, we developed a murine experimental system to manipulate proprioceptive inputs during memory-dependent localization task (which required precise operant-conditioned licks) in spatial virtual reality (VR). Through bioluminescent optogenetics (luminopsin) selectively expressed in the parvalbumin-positive neurons of the dorsal root ganglia in mice, proprioceptive processing was compromised directly from the periphery to bypass the bottleneck of specific central targeting, which results from the lack of anatomically or genetically dedicated proprioceptive circuits in the brain. *In-vivo* IVIS imaging and behavior suggested the effects of luminopsin last for roughly 20 minutes. While mice exhibited normal performance in tasks relying on gross motor skills, they showed subtle deficits in challenging spatial tasks that required integration of past movements. These observations support a task-specific role for proprioception, and demonstrate a potential of chemogenetics-like, rapidly reversible strategies for characterizing peripherally defined sensory contribution to spatial cognition. Future work will optimize this approach; for instance, to activate opsins by light with millisecond precision. To our knowledge, this is a first causal demonstration for acute participation of proprioception in path-integration biomechanics, enabling the first temporally defined method for mild perturbation of path-integration mechanisms.

## Introduction

Spatial navigation requires integration of external sensory inputs and internal generative and feedback signals, allowing animals to construct cognitive representations of their environments and execute goal-directed movements^1^. The central nervous system (CNS) processes these inputs within two major computational frameworks: allocentric and egocentric reference frames^2^. Allocentric navigation is concerned with spatial relationships relatively independently of the observer, whereas egocentric navigation considers spatial information primarily relative to the individual. Path integration, a core aspect of egocentric navigation, may be pivotal to use of the information in both frameworks, where self-motion (idiothetic) cues update neural representations of the position even in the absence of external landmarks (i.e., allothetic cues)^3^.

Among self-motion cues, proprioception is historically difficult to investigate regarding its causal functional role and the underlying neural basis. As a result, data for causal, quantitative contribution of path-integration biomechanics to navigation are scarce. As perception for muscle and joint position and kinematics derived from mechanosensory feedback ^4,5^, proprioception is crucial for motor control, yet its contribution to spatial cognition remains unclear. Traditionally, studies on path integration have focused on vestibular (especially head-direction)^6,7^ input, with specific work done for proprioception very limited. Nevertheless, observational evidence implied that proprioception had a contribution to real- or virtual-world navigation, including orienting tasks and spatial memory recalls of different reference frames ^8–12^. While the hippocampus and entorhinal cortex are recognized as key centers for spatial navigation, how proprioceptive signals influence the processes in these higher cognitive areas is largely unknown, at least not in a specific and causal sense (see ^13–15^ for elegant model-driven work and observations). There are urgent needs to find a strategy for mild, reversible manipulation of proprioceptive processing to address the scientific barriers.

Neurobiological research has identified distinct neural pathways for conscious and subconscious proprioceptive processing^5,16–18^. Dorsal root ganglion (DRG) neurons transmit proprioceptive signals via Aα and Aβ fibers, which project to the somatosensory cortex^19,20^ and cerebellum. Conscious proprioceptive pathways contribute to fine motor control and voluntary movement, whereas subconscious pathways, including the spinocerebellar tract, enable postural adjustments^16,17,21^. A continually updating view for proprioception is its potential contribution to higher cognitive processes, such as emerging evidence for the cerebellum to shape the hippocampal spatial code. This is consistent with the notion that proprioception is supporting conscious and subconscious body perception^40^. Although relevant mechanisms of lower level control and sensation are well characterized, how proprioceptive information interacts with navigation-relevant circuits to shape spatial navigation^22^ remains speculative with little interventional evidence, like in animal models..

A major challenge in establishing the causal role of proprioception, as well as biomechanics-based path integration, in spatial navigation is the difficulty in selectively perturbing proprioceptive signals without affecting other sensory or motor functions. Existing genetic knockout approaches, such as *Piezo2* deletion^23,24^, often result in severe motor impairments. Moreover, conventional methods targeting proprioceptive pathways lack the temporal control, to avoid biological compensation or learning alternative neuro-behavioral strategies.

To address these limitations, we developed an experimental framework that combines virtual reality (VR) with bioluminescent optogenetics to selectively manipulate proprioceptive feedback in behaving mice. The choice of head-restrained VR will confine the utility of self-motion components as a pragmatic condition for identifying the computational role of proprioception^22,25^. Using promoter-dependent transgenics of Luminopsin 3 (LMO3), an opsin-luciferase fusion protein^26^, we transiently modulated parvalbumin (PV)-expressing proprioceptor DRG neurons^27^ via local administration of coelenterazine (CTZ) to the muscles of the limbs. We expect that luminopsin activation depolarizes PV^+^ DRG proprioceptors and induces aberrant firing, thereby introducing corruptive proprioceptive signals that mismatch the actual muscle state, as PV-dependent interference has been shown to impair precise proprioceptive functions^27^. This spatially local, temporally restricted and genetically targeting strategy created a time window of ∼20 minutes for temporary and quickly reversible perturbation of normal proprioceptive signaling, directly at the periphery where the modality-specific neural pathway is still isolated^19,28^. As an outcome, this approach interfered with proprioception selectively without disruptions of motor functions and possibly peri-personal sensation^20^ as in conventional knockouts.

Two complementary lines of arguments were employed to establish an experimental framework for evaluating whether and how acute proprioceptive perturbation affected memory-dependent spatial computation. First, the simplified neuroethological environment we created provided a highly quantitative and interpretable paradigm for subtle effects. We hypothesized that mice with impaired proprioceptive feedback would exhibit altered performance in tasks requiring the integration of past movements to locate a virtual goal. By a closed-loop VR, it enabled an assessment of how non-acceleration self-motion signals contribute to spatial decision-making. Second, having limited the effects of perturbation by strict ligand delivery to local muscles, we still conducted a series of control experiments to support the restricted scope of PV-dependent localized manipulation. Our results suggested that proprioceptive modulation selectively influenced specific aspects of navigation-related responses, while leaving general locomotor abilities and other motivational and simple coordinated behaviors intact. This study points toward a possible new insight into the task-specific role of proprioception in spatial cognition, and demonstrates the utility of chemogenetics manipulation for probing sensorimotor integration for proprioception. This may as well be a first temporally defined neural manipulation for path integration in the computation of spatial estimates in rodents.

## Results

Our overall goal was to establish a novel method that could produce transient, reversible perturbation selectively to the proprioception, to the extent just sufficient to affect computation of self-motion cues for navigation without “leaking” into other domains, like mobility^23^, motor control^27^, balance, motivation, or even sngception^29^ or induction of chronic pain. A sensitive, quantitative and reliable paradigm is necessary to detect the subtle changes this perturbation generates over the computation of navigation-related signals, and our choice was a head-restrained design of fictive navigation in VR.

### Considerations for using transgenics with the bioluminescent optogenetic actuator LMO3 in proprioceptive neurons of the DRG

To begin, we carried out a series of work for considering the reliability of the transgenics, the dosage and the location of administrating the ligand for luminopsins. To validate LMO3 expression in our target DRG PV^+^ neurons, we extracted the DRG associated with the 3^rd^ lumbar vertebra (L3) of the spinal column, which receives somatosensory input from the largest muscle group, the *quadriceps femoris*^30^, in the PV-Cre;td-LMO3 mice. Fluorescent immunocytochemistry was conducted to label LMO3 channels (Fig. **1A, B**). LMO3^+^ cells and PV^+^ cells were highly colocalized, with 66% of PV^+^ neurons expressing the LMO3 proteins; on the other hand, the most majority (86%) of LMO3 proteins were expressed in the PV^+^ neurons (Fig. **1C**; n = 3 mice). LMO3 channels were not expressed in muscles because of the *synapsin* promoter used for the LMO3 locus. Although the LMO3 expression pattern displayed some lever of aggregation (Fig. 1B, *green*), it was consistent with the observations reported in the original publication and deemed functional based on the experiments^26,31,32^; see also ^27^).

**Figure 1.**
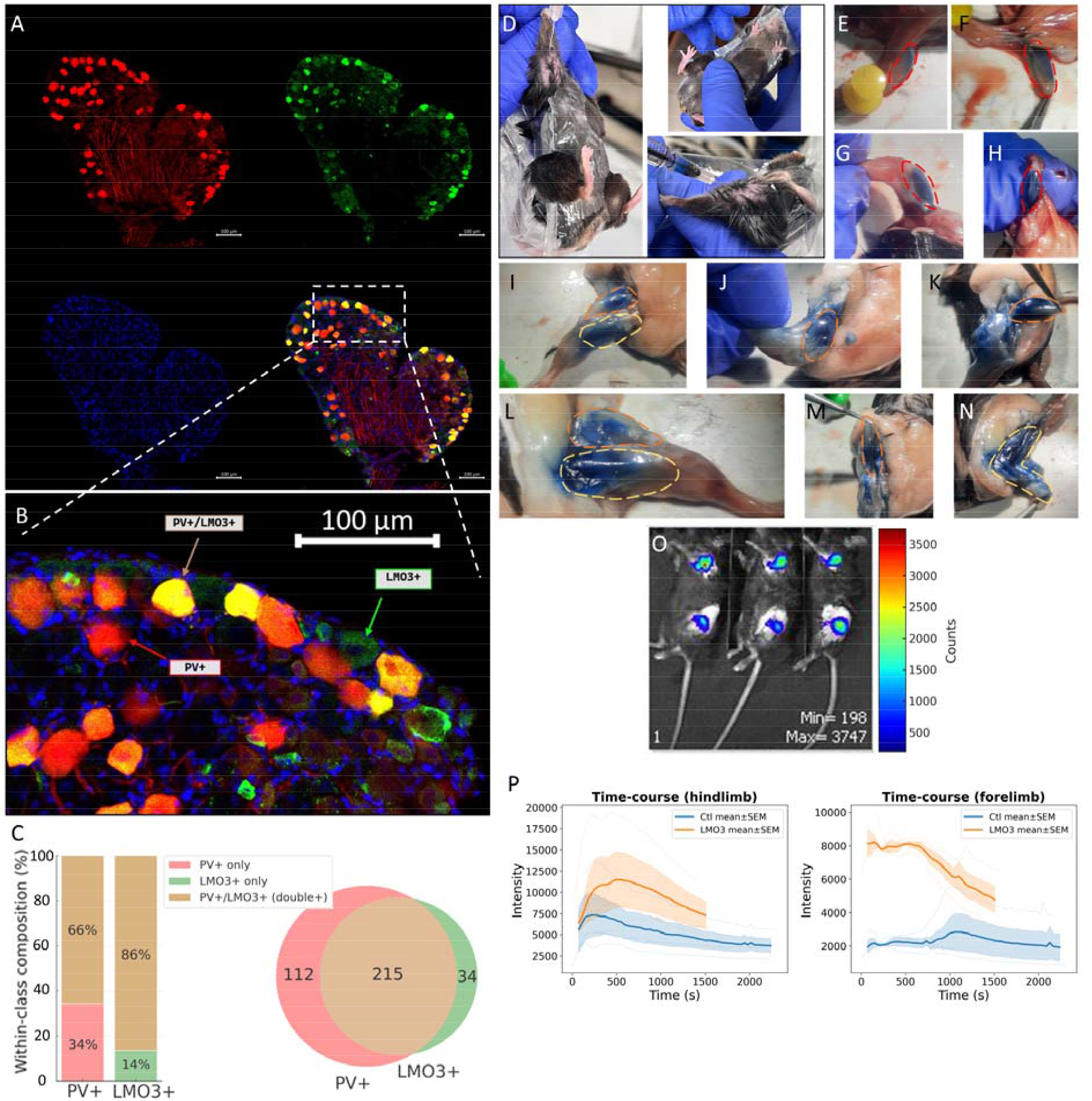
Characterization of LMO3 Expression in Proprioceptive DRG Neurons, Validation of Intramuscular Injection Sites, and Assessment of Bioluminescence Following CTZ Administration (A, B) Protein expression patterns of LMO3 and Parvalbumin (PV) in dorsal root ganglia (DRG). Representative confocal images of a whole-mount DRG. Shown are tdTomato-expressing PV⁺ neurons (red, top left), LMO3⁺ neurons labeled by immunostaining (green, top right), and cell nuclei stained with DAPI (blue, bottom left). The merged image (bottom right) indicates co-localization (yellow) of PV and LMO3. Scale bars = 100 µm. **(C)** Quantification of neuronal populations. The bar chart (left) shows that 66% of PV⁺ neurons co-express LMO3, and 86% of LMO3⁺ neurons co-express PV, based on data from n = 3 mice. The Venn diagram (right) depicts the absolute counts and overlap between the PV⁺ and LMO3⁺ populations. **(D-N)** Anatomical validation of intramuscular injection coverage. **(D)** Demonstration of the intramuscular injection technique in a restrained mouse. **(E-H)** Post-mortem dissection images showing the spread of 2% Evans blue dye within the triceps brachii of the forelimb (outlined in red) after injection. **(I-N)** Dissection images confirming dye coverage in the hindlimb muscles, including the quadriceps femoris (outlined in orange) and hamstring muscles (outlined in yellow). **(O-P)** *In-vivo* bioluminescence imaging (IVIS) after intramuscular CTZ delivery. **(O)** Representative IVIS images of three mice showing bioluminescence signals in the forelimb and hindlimb regions following injections. **(P)** Graph of bioluminescent signal intensity (photo count per second) over time for the forelimb and hindlimb injection sites. Bioluminescence is expressed as photon counts per second per square centimeter. Note that signal traces were aligned according to the time of injection (points for baseline signals were included before the injection time), and the rise phase of bioluminescence was mostly omitted; only one mouse (thin lines indicate data from individual animals) which received injections as the last of the batch imaging procedure had data recorded right after the injection time. Data are presented as mean ± SEM. Mixed-design linear model regression.

Multiple tests have been performed to determine the site of drug administration and their dosages. Considering the convenience of administration, we tried intraperitoneal (*i.p.*) injection of CTZ **(20 mg/kg body weight)** yet observed no clear effects in the head-fixed dead reckoning test (see **Fig. S5**). We generally worried that CTZ went through the blood-brain barrier and confounded the result interpretation by also affecting PV^+^ cells in the brain, so used local intramuscular (*i.m.*) injection instead. Initially, we targeted the muscles of the distal limbs (ventral forearm muscles and *gastrocnemius*) using a relatively low and medium CTZ dosage **(0.1 and 2.5 mg/kg body weight)**^29^. However, preliminary behavioral experiments showed no significant result for the effects of the CTZ **(Fig. S3, S4)**. This led us to reassess the target muscle and CTZ dosage for our specific experimental objectives and design. As opposed to focusing on induction of sng (muscle soreness) and chronic pain, which presumably required only a transient, priming-like proprioceptor stimulation with a low-dose treatment of CTZ, our study aimed to investigate behaviors and computations related to ongoing muscle mechanical feedback, which would necessitate a more sustained modulation.

To assess whether the lack of positive results was due to relatively indirect interference on locomotion resulting from perturbed limb functions, we switched the injection target to proximal limb muscles (*Triceps brachii*, *Quadriceps femoris* and *Hamstring*) which is supposed to be a primary drive of the stepping kinetics.

Regarding CTZ dosage, our goal was to achieve sustained modulation, leading to an increase of the dosage. However, we noted the absence of published work done with CTZ applied to manipulate excitability of peripheral neurons. Consequently, we referenced studies on luminopsin tool development and peripheral nerve regeneration^33,34^ and increased the dosage to **8.2 mg/kg body weight**.

To accommodate the short period of effective time for luminopsin activation, we developed and optimized a new method for injecting the ligand without a need of anesthetizing the mouse and waiting for its recovery. In our pilot experiments, the effect of isoflurane anesthesia led to apparent reduced of the mouse activity in the head-restrained VR for the first 10-30 minutes (data not shown). We replaced gaseous anesthesia with a body-constraint method: a multi-holed, funnel-shaped plastic bag was used to secure the animal for intramuscle injections. When this technique was performed skillfully, struggling of the mouse as well as post-injection low mobility were minimized.

To validate the spatial accuracy of our injection method, 2% Evans Blue was injected to the target muscles for examination. As in **Fig. 1D-N**, this technique allowed for reasonable muscle targeting even in conscious, un-anaesthetized animals. We could see *Triceps brachii*, *Quadriceps femoris* and *Hamstring* were clearly and selectively stained in blue.

The duration (or *in-vivo* lifetime) of CTZ effects posed another challenge, as no existing studies we knew had systematically explored CTZ’s impact during animal behavior. From our best estimates from references^33^, anecdotal observations and personal communications, research involving use of CTZ for other purposes (including LMO tool development, peripheral nerve regeneration and chronic pain) suggested that its effects could begin around **30 seconds** after IP injection and decay sometime within **1 hour**. To obtain empirically based estimates for the physiological effects of luminopsin following a single injection of CTZ, we performed *in-vivo* IVIS imaging on anesthetized mice with the injection of the given location and dose. In this set of experiments, bioluminescence of LMO3 activation was detected as a proxy of LMO3 functions, and the PV-Cre;LMO3^f/f^ mice showed ∼20 minutes of significant stronger detectable signals in both hindlimb and forelimb (n = 3 mice; **Fig. 1O, P**), in contrast to the control (n = 3 mice). These timeframes could primarily be influenced by pharmacodynamics for the presence of chemical in the muscles given that the activation/deactivation of luminopsin by CTZ is much faster^35^. Importantly, the result of our later behavioral experiment using walking on the metal grid (see **Fig. 5**) was consistent with this time-scale of the injection effect, in addition to consistency with a similar IVIS assessment of the bioluminescence from a different system for mice (particularly at a similar 5 mg/kg body weight dosage)^34^.

### Normal Locomotor Activity, 2-D Motion Kinematics, Anxiety and Motivation for Reward Seeking in Mice with LMO3-Perturbed Proprioceptive Nerves in Limbs

Manipulation of mechanical sensation may impair an animal’s ability to perform locomotion or its motivation to move, as shown by *piezo* knockout, where mice exhibited significant difficulties in normal movement^23,24^. To test whether the LMO3 system produced similar impairments, we conducted two behavioral tests, Open Field and Rotarod, and additionally used Lever Pressing in a Skinner-box setting to assess reward-seeking behaviour even when cumulative efforts are required^36,37^. These tests were designed to evaluate basic locomotion and proxy for the anxiety level, motor coordination, and motivation for acquiring rewards, respectively. For the Open Field, we used a between-subjects design: opsin-expressing PV-Cre;LMO3^f/f^ mice were compared with pooled opsin-negative controls (wild-type and PV-Cre), with all animals receiving a high dose of CTZ (8.2 mg/kg body weight; 180 µg in total) delivered to the proximal limb muscles(5 µL to the forelimbs, 20 µL to the hindlimbs), so that a group difference would reflect LMO3-mediated proprioceptive perturbation rather than CTZ injection itself. For the Rotarod and effort-based Lever Pressing, saline injection served as a within-subject pharmacological control. All mice were of the PV-Cre;LMO3^f/f^ genotype, receiving CTZ at 0.1 mg/kg to the distal limbs (with the same total volumes).

In the Open Field, the freely moving behaviors of mice was characterized in a 2-D arena over 30 minutes. Here we compared opsin-expressing PV-Cre;LMO3^f/f^ mice with pooled opsin-negative controls (wild-type and PV-Cre), so that a difference would isolate LMO3-mediated proprioceptive perturbation from the effect of CTZ injection itself. Across the traditional kinematic measures, PV-Cre;LMO3^f/f^ mice showed a non-significant trend toward more locomotion (**Fig. 2A-D**) — and more time in the central zone (**Fig. 2E**); however, none of these small differences survived correction (all q ≥ 0.119). The pooled occupancy maps were near-identical, both dominated by thigmotaxis (**Fig. 2F, G**). Per-frame velocity and acceleration vector distributions were indistinguishable (**Fig. 2H, I**; energy-distance permutation, q = 0.068). Using a novel, training-free hierarchical behavioral clustering and unsupervised discovery method^57^, we identified 9 reliable behavioral classes during navigation, validated by the region-of-interest (ROI) of raw images. Among them, the only class difference survived multiple-comparison correction was the time spent turning left (Δ = 0.023, 95% CI [0.011, 0.034], q = 0.027). No other differences were found in behavioral classes between the control and PV perturbation group (**Fig. 2J, 2K**). Overall, we think that mice with localized CTZ injections, which induced a transient, priming-like proprioceptive stimulation in limbs, showed broad invariance of free 2-D navigation and anxiety metrics across traditional parametric and unbiased machine-learning analyses. The effects were restricted without altering general behavioral capacity.

**Figure 2.**
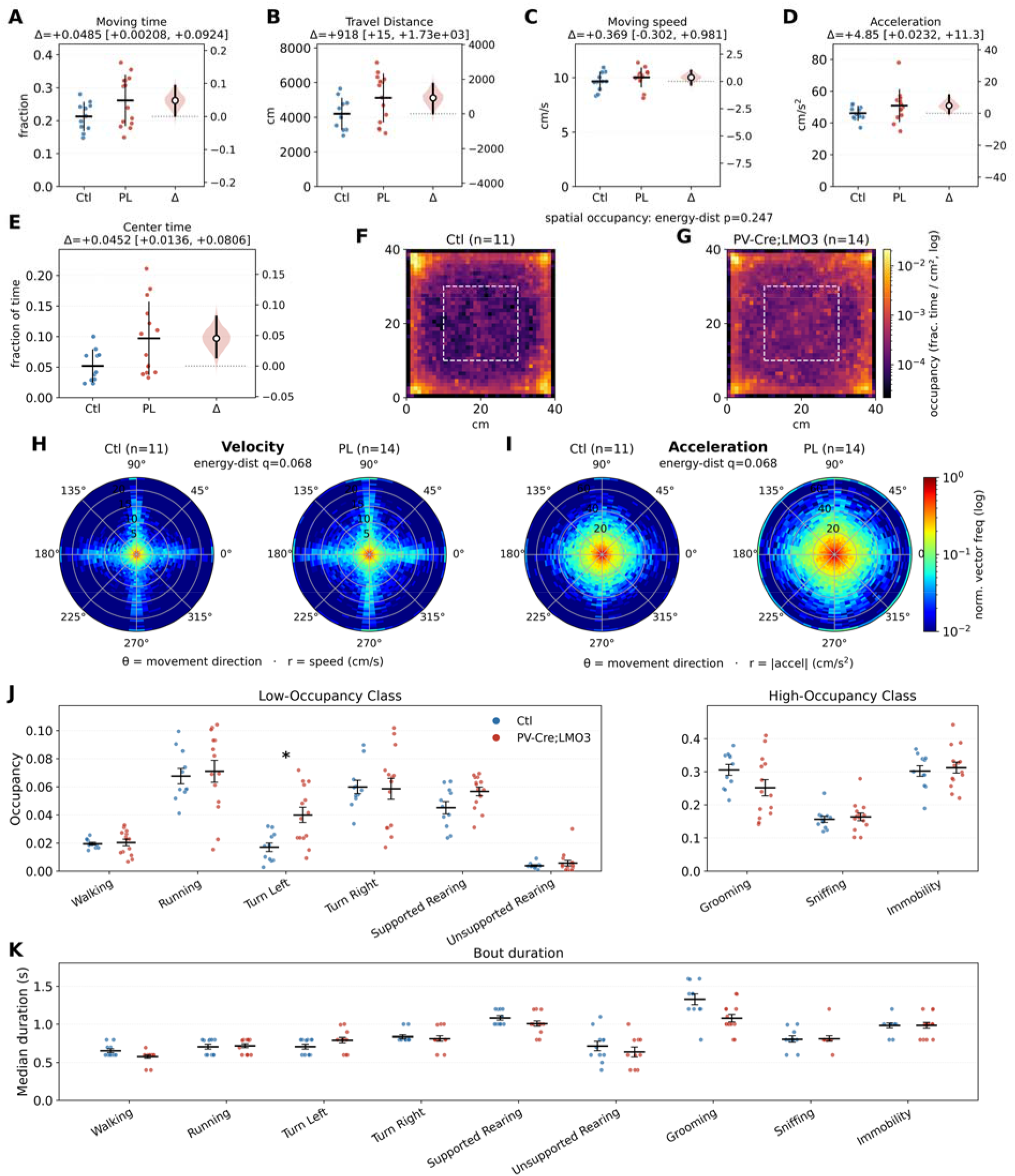
Mice with intramuscularly injected CTZ showed no consistent changes in 2-D curvilinear motion kinematics, open-space anxiety proxy as well as gross exploratory behavioral classes in an Open Field arena. Opsin-expressing PV-Cre;LMO3^f/f^ mice (PL, *n* = 14) were compared with opsin-negative controls (wild-type and PV-Cre; Ctl, *n* = 11), both receiving matched proximal high-dose CTZ (5 μL for the forelimbs and 20 μL for the hindlimbs; 8.2 mg/kg body weight). All measures were calculated from the post-injection 5–30 min time window. Significance was tested by unpaired permutation test (5,000 resamples) and corrected for multiple comparisons with the Benjamini–Hochberg false-discovery-rate. \**q* < 0.05. **(A–E)** Locomotor kinematics and proxy of behavioral anxiety, shown as Gardner–Altman estimation plots (raw groups are summarized as mean ± s.d. to depict dispersion; statistical inference is given by the bootstrap 95% CI of the mean difference and the permutation test^56^): fraction of time moving (A), distance travelled (B), mean speed while moving (C), mean absolute acceleration while moving (D), and fraction of time in the central zone (the central 20 × 20 cm area) (E). In each panel, raw per-animal values are plotted at left (horizontal bar, mean; vertical bar, s.d.) against the mean-difference (Δ), the observed effect size, at right (shaded curve, bootstrap distribution of Δ; open circle, Δ; vertical line, 95% CI). Shadowed curves show a resampled distribution of Δ based on observed data (bootstrap); CI, 95% bias-corrected and accelerated (BCa) bootstrap confidence-interval of resampled Δ. **(F, G)** Group-mean spatial-occupancy map for Ctl (F) and PL (G) shown in log scale (dashed square, central zone). Both groups displayed pronounced thigmotaxis. Animal-level energy-distance permutation test (1,000 resamples), p = 0.247. **(H, I)** Group-mean radial heat map for polar representation of per-step (based on down-sampled time bins of 200 ms) velocity (H) and acceleration (I) vector distribution: angular position, movement direction; radius, vector magnitude (capped at the 99.5th percentile); color, panel-specific normalized frequency (log scale). *q*, animal-level energy-distance permutation test (1,000 resamples). **(J)** Fractional occupancy of behavior class, shown according to low- and high-occupancy classes for visualization. Unpaired permutation test of the mean difference (5,000 resamples); asterisks denote q < 0.05 after Benjamini–Hochberg false-discovery-rate correction applied across all nine occupancy classes. **(K)** Median per-bout duration of each behavior class. All whisker representations in J and K are mean ± s.e.m. q > 0.05 across groups, unpaired permutation test of the mean difference (5,000 resamples) with Benjamini–Hochberg false-discovery-rate correction applied.

In the Rotarod test, mice were tested for their gross motor coordination of limb movement and body balance on an accelerating rod (**Fig. 3B**). There were 3 test sessions, and the order was arranged in a way to counter-balance the animal’s experience (as the learning effects observed with repetition were obvious), and also show the experience-dependent effect (comparing two consecutive sessions with the same treatment) (**Fig. 3A**). There was no significant difference in the time spent on the rod between PV-Cre;LMO3^f/f^ mice that received saline and CTZ (**Fig. 3C**).

**Figure 3.**
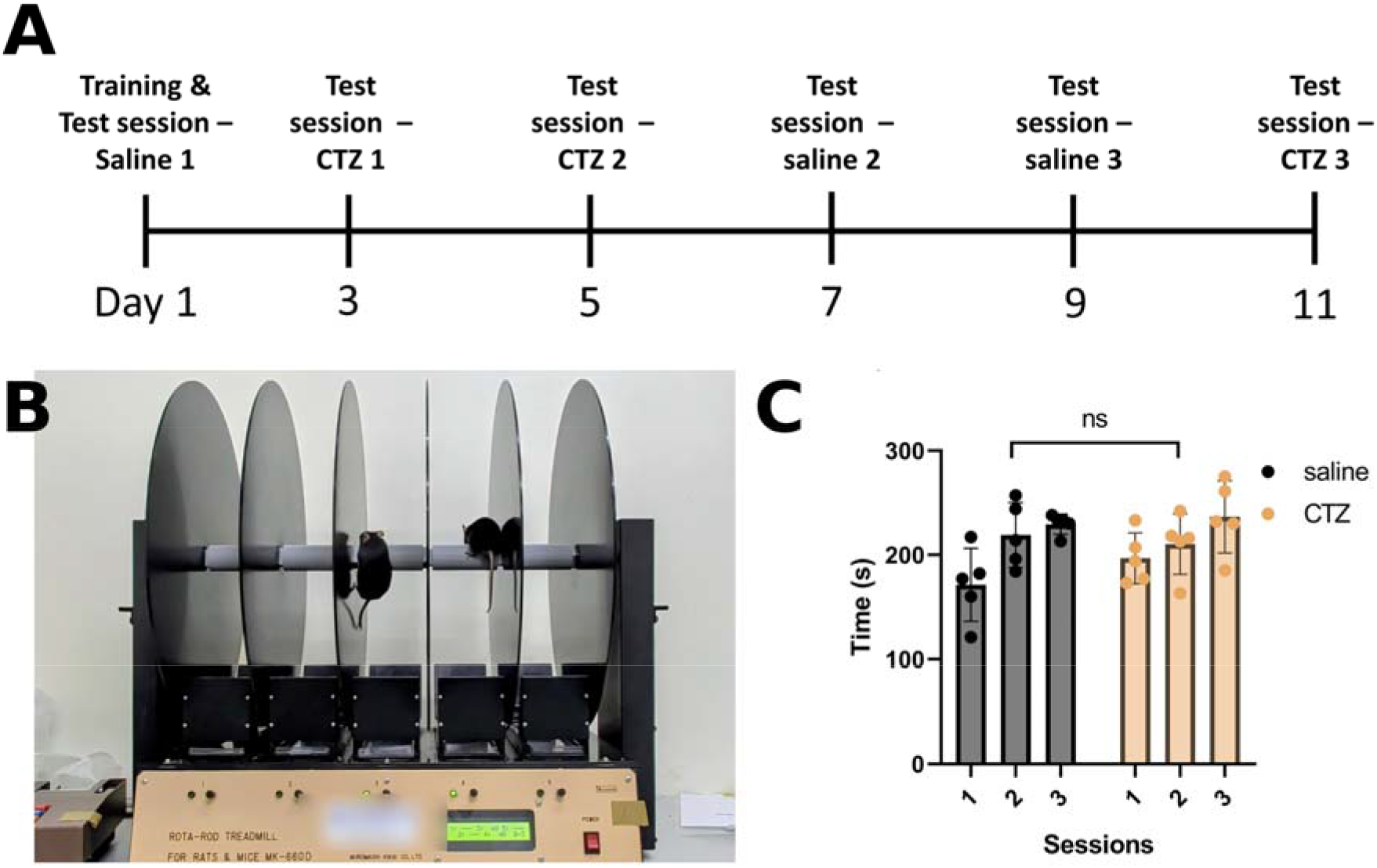
Mice with intramuscularly injected CTZ showed no difference in simple motor coordination and muscle endurance on a rotarod. This experiment used the treatment as the control, and all mice were of the PV-Cre;LMO3^f/f^ genotype. The dose of CTZ was 0.1 mg/kg, with a total injection volume of 5 μL for the forelimbs and 20 μL for the hindlimbs, administered to the distal limbs. **(A)** Experimental schedule of the rotarod test. The schedule utilized an interleaved design to minimize the impact of learning experiences. **(B)** Photograph of the rotarod (accelerating-rod) apparatus. **(C)** Comparison of the time stay on the rod. Data represent mean ± SEM. Saline n = 5 mice, CTZ n = 5 mice; repeated measures two-way ANOVA; no significant difference; n.s.

Finally, the effort-based Lever-Press test was designed to evaluate motivation for reward-seeking behavior. With a custom-designed Skinner Box setup (Fig. **4A**), a structured training protocol was applied to ensure that mice learned the association between lever-pressing and receiving sucrose solution as a reward (**Fig. 4B**; see **Methods)**. In the Progressive Ratio (PR) phase, the effort required for successive rewards was designed to follow an exponential growth (**Fig. 4C**). This task structure was made for detecting a true level of efforts the animal was willing to spend while the reward-seeking regime was kept well below water-satiety. There were no significant differences found between PV-Cre;LMO3^f/f^ mice and their genetically matched controls, with both groups receiving CTZ injections (**Fig. 4D, E**), indicating no difference in reward-associated motivation by perturbing PV^+^ neurons with injections made to the muscles.

**Figure 4.**
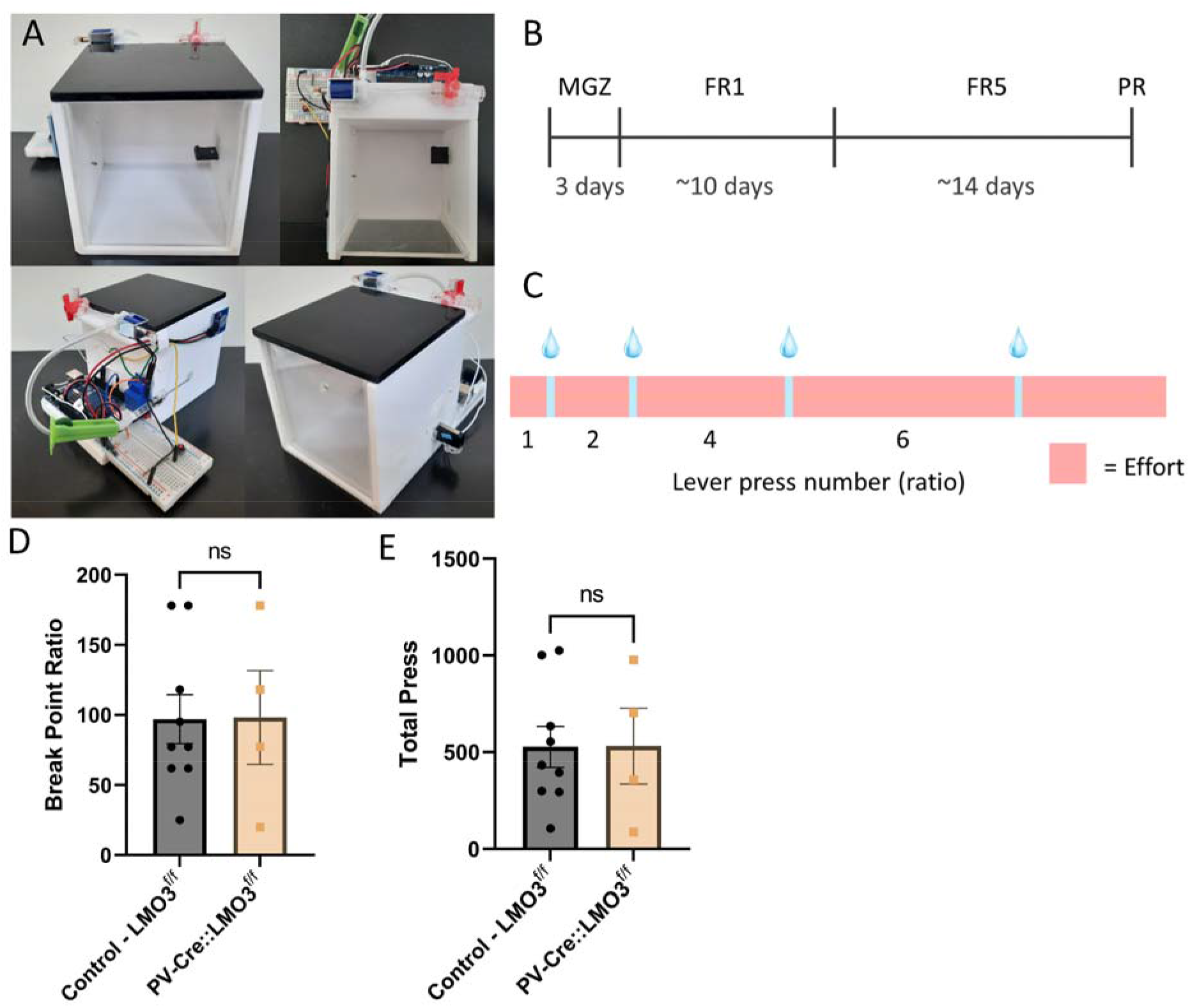
Mice with intramuscularly injected CTZ showed no difference in reward-seeking behavior with effort-based, progressive ratio (PR) task design. The dose of CTZ was 0.1 mg/kg, with a total injection volume of 5 μL for the forelimbs and 20 μL for the hindlimbs, administered to the distal limbs. **(A)** Actual appearance of the custom-designed Skinner box. **(B)** Experiment schedule. Magazine Training (MGZ), Fixed Ratio (FR), Progressive Ratio (PR). **(C)** Effort-based progressive ratio design schematic. **(D, E)** Comparison of the metrics implying the efforts mice paid. Data represent mean ± SEM. Ctl (LMO3^f/f^) n = 9 mice, PV-Cre;LMO3^f/f^ n = 4 mice; Wilcoxon rank-sum test; no significant difference; n.s.

We understand the limitation of the Rotarod and Lever-Pressing experiments as using possibly less effective injection dose and sites to optimally interfere with certain task-relevant proprioception. On observations across all tasks, however, there was no reason to think the higher dose and injection site (proximal limbs) affected mice’s general behaviors, but quantitative assessments could be further improved. Due to scheduling and resource constraints, we did not repeat all of these tests after testing over a range of injection locations and dosages. Although our experiments described below support a specific manipulation made with local injections at a high dose of CTZ, these results serve to confirm that the restraint, injections and proper luminopsin activation did not incapacitate the animals for using their general faculty to perform motion-based, goal-directed tasks.

### Locomotor coordination and proprioceptive feedback showed limited and selective impairment under LMO3 perturbation

To assess whether the LMO3 system affects motor coordination, we performed two locomotion-related tasks^27^: Grid Walking and Balance Beam. The Grid-Walking task evaluates free locomotion on discrete metal grid (of grid thickness 1.5 mm and spacing 18.5 mm) that requires fine mechanical feedback by depriving the animal of visual (in darkness) and whisker (trimmed; see Methods) sensory inputs (**Fig. 5A**). Notably, the travel path of each animal was tracked for normalizing foot fault count according to each individual’s mobility. In the standard protocol, animals performed the task for 5 minutes. Under low-dose CTZ (0.1 mg/kg body weight) injected into the distal limb muscles, no significant differences were observed (**Fig. 5B**). We further extended the task duration to 30 minutes to address the time-dependency of the CTZ effects. The behavior was analyzed with a temporal binning of 5 minutes, and foot faults were quantified separately for each limb (see Methods for details). With the high CTZ dose (8.2 mg/kg body weight) administered to proximal limb muscles, the PV-LMO3 animals exhibited significantly more foot faults from the hindlimb compared to controls (Ctl) during the 5–30 minute period (Ctl (LMO3^f/f^): n = 12 mice, PV-Cre;LMO3^f/f^: n = 7 mice; ART-corrected ANOVA test, p = 0.0231; **Fig. 5D**). A more careful visualization and resampling method revealed the effect size, which can be more informative the statistical significance (Fig. **5C**; see Methods). Although *post-hoc* analyses of individual time bins did not reveal statistically significant differences **(Fig. S2)**, it appeared that the most significant time bin fell within the first 10 minutes, qualitatively consistent with our estimation of the temporal kinetics of CTZ effects with IVIS imaging (c.f. **Fig. 1**).

**Figure 5.**
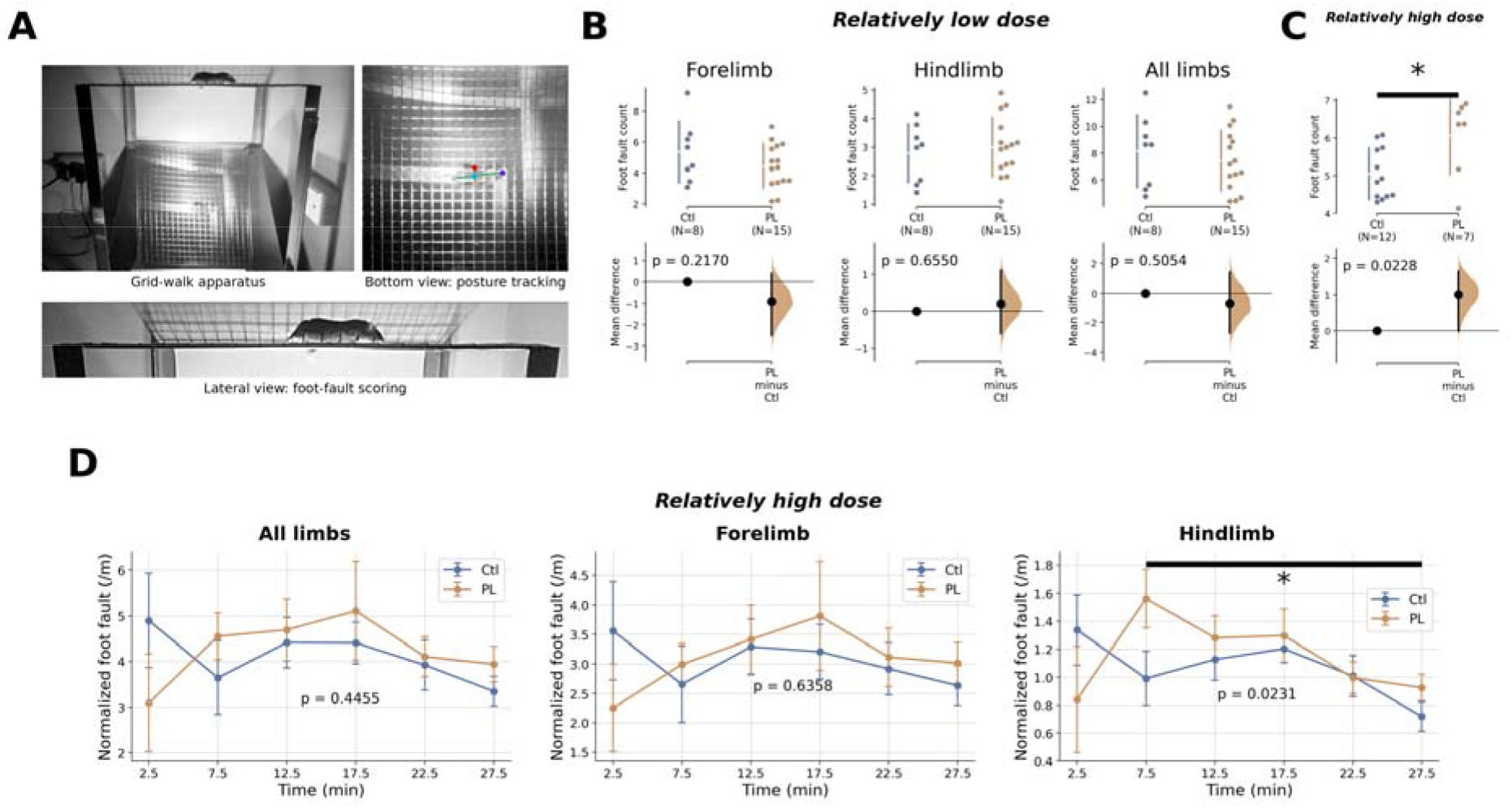
Whisker-trimmed mice with intramuscularly injected CTZ showed significant but subtle difference in foot fault when walking on a discrete metal grid in darkness. In **(B)**, mice received distal muscle injections of CTZ at a **relatively low dose (0.1 mg/kg body weight)**. In **(C, D)**, mice received proximal muscle injections of CTZ at a **relatively high dose (8.2 mg/kg body weight)**. Cumming estimation plots in **(B) and (C)** illustrate the mean and ± standard deviation for each group, represented by the gap and vertical ends of gapped lines. Mean difference (Δ) displays the effect size. Shadowed curve indicates a resampled distribution of Δ based on observed data (bootstrap), with the black filled circle and vertical line depicting the mean and 95% confidence interval of resampled Δ. Due to post-handling neuro-behavioral instability, statistical analysis was performed only over 5-30 mins in (**D**). **(A)** Photographs of the Grid Walking task in complete darkness. Lateral view for foot fault scoring; bottom view for DLC-tracked posture and locomotion kinematics, all taken under infrared videography. **(B)** There is not significant across comparison at the relatively lower dose. Ctl (LMO3^f/f^) n = 8 mice, PV-Cre;LMO3^f/f^ (PL) n = 15 mice. Unpaired permutation test. **(C)** Significantly more foot fault at the relatively higher dose. Ctl (LMO3^f/f^) n = 12 mice, PV-Cre;LMO3^f/f^ (PL) n = 7 mice. Unpaired permutation test; *p < 0.05. **(D)** Time- and injection-site-dependent effect on foot faults is consistent with the spatial and temporal specificity of luminopsin in mice’s fine motor coordination. Point and lines indicate mean ± s.e.m. Ctl (LMO3^f/f^) n = 12 mice, PV-Cre;LMO3^f/f^ (PL) n = 7 mice. ART-corrected ANOVA test; *p < 0.05.

The task of Balance Beam is a stress-driven assay for fine motor coordination (**Fig. 6A**). Under the elevated condition and unpleasant light illumination set up at the start point of the beam, the coordination of hindlimbs (which the animal could not see) with overall motor actions is particularly needed during walking over the narrow beam. Previous study demonstrated that PV-Cre;ASIC3^f/f^ mice exhibited significantly more foot faults, suggesting impaired proprioception^27^. In contrast, in our study, PV-Cre;LMO3^f/f^ mice with CTZ injections at both the low doses (0.1 mg/kg body weight, distal limb muscles; **Fig. 6B-E**) and the **high dose** (8.2 mg/kg body weight, proximal limb muscles; **Fig. 6F-H**) showed no significant differences in travel time or the number of hind-paw foot fault compared to controls, across different widths of the beam used (see **Method Details**). The experiments were carefully designed to examine both *within*- and *between*-subject effects. Notably, we have not stress-tested the animals with injecting CTZ to the limbs unilaterally, as suggested in the earlier work. To sum up, different tasks showed different sensitivity to the same luminopsin perturbation system, which would suggest a task-context-dependent computation.

**Figure 6.**
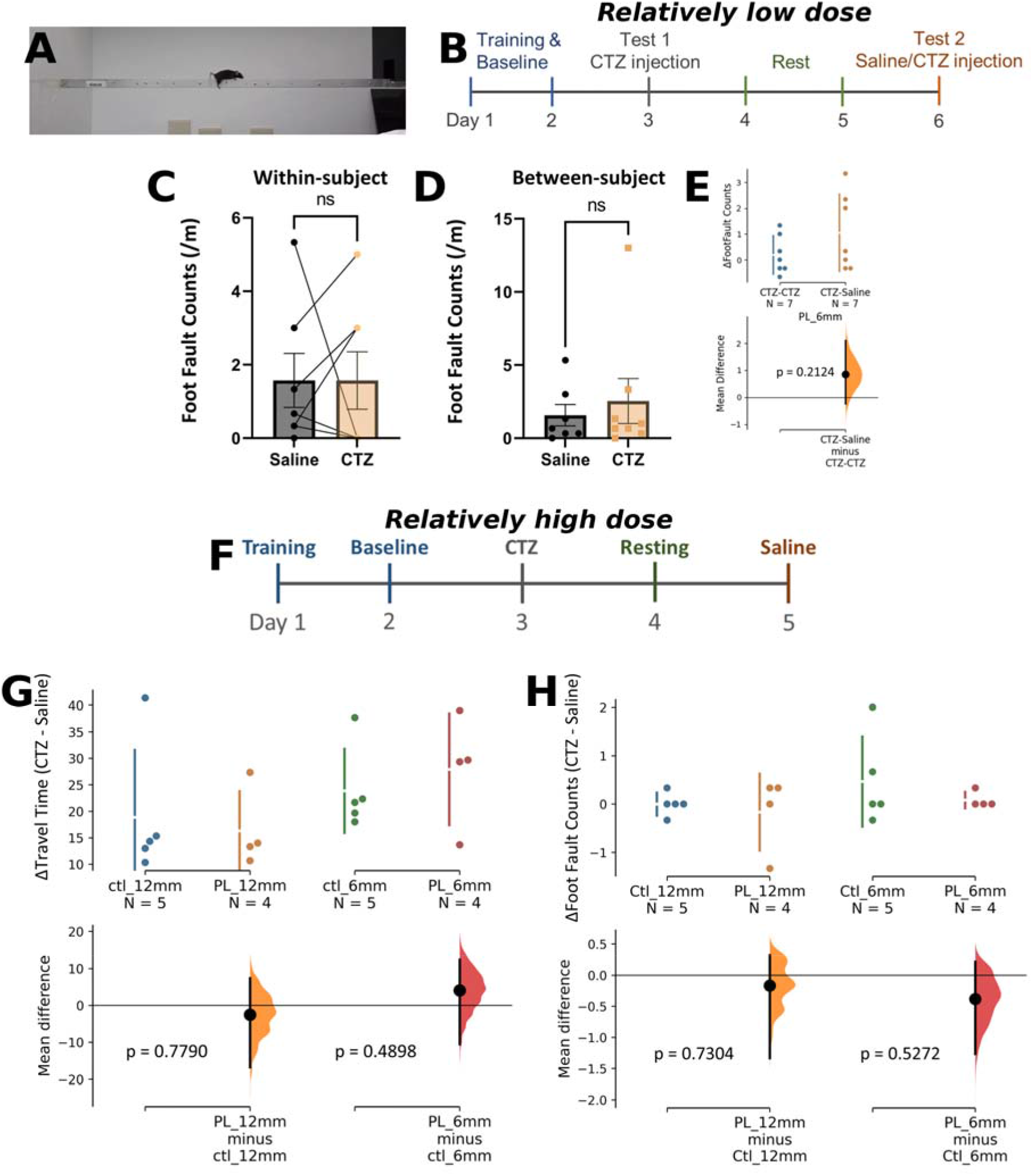
Mice with intramuscularly injected CTZ showed no apparent difference in fine motor coordination on a balance beam. The Cumming estimation plots (E, G, H) are similar in format to those in Figure 5. In (C-E), mice received distal muscle injections of CTZ at a relatively low dose (0.1 mg/kg body weight). Ctl (LMO3^f/f^) n = 7 mice, PV-Cre;LMO3^f/f^ (PL) n = 7 mice. In (G, H), mice received proximal muscle injections of CTZ at a relatively high dose (8.2 mg/kg body weight). Ctl n = 5 mice, PL n = 4 mice. no significant difference; n.s. **(A)** Photograph of the balance beam apparatus (a mouse traversing the 6-mm beam). **(B)** Experimental schedule for the balance beam test with a relatively low dose distal limb injection. **(C)** Foot fault count comparison within genetic group. Paired Wilcoxon rank-sum test. **(D)** Foot fault count comparison cross genetic group. Wilcoxon rank-sum test. **(E)** Cumming estimation plot comparing Δfoot faults. Unpaired permutation test. **(F)** Experimental schedule for the balance beam test with a relatively high dose proximal limb injection. **(G)** Cumming estimation plot comparing Δtravel time. permutation test. **(H)** Cumming estimation plot comparing Δfoot faults. permutation test.

### Design of a VR-like, self-motion-based system for sensitive and quantifiable spatial anticipatory behavior

The transient luminopsin-based perturbation may be subtle to mice, which is an important feature we need to mildly manipulate the proprioceptive information flow, without affecting the gross and overall sensorimotor faculties during goal-directed navigation. To further confirm selective degradation of proprioceptive function in path integration, a core basis of navigation, it is crucial to have a behavioral system highly sensitive and quantitative for readout of that. We set out to design a system inspired by our animal VR research which measured subtle spatially dependent behavior in anticipation of a memorized location (see **Supplemental Method Details**).

In this task of our design, every head-fixed mouse navigated through a linear virtual environment (**Fig. 7A**). The environment featured a starting auditory cue within the first 0–5 cm, followed by no external cues throughout the same trial. Upon entering the RZ and licking, the animal received a drop of sucrose water as a reward. After receiving the reward or reaching the end of the trial (whichever happened first), the system transitioned to an inter-trial interval (ITI) state. During the ITI, if the animal ceased movement, it was teleported back to the starting point for the beginning of the next trial (**Fig. 7B**; see **Fig. 7C** for electro-mechanical integration with *Bonsai*, and **Supplemental Method Details** for more information).

**Figure 7.**
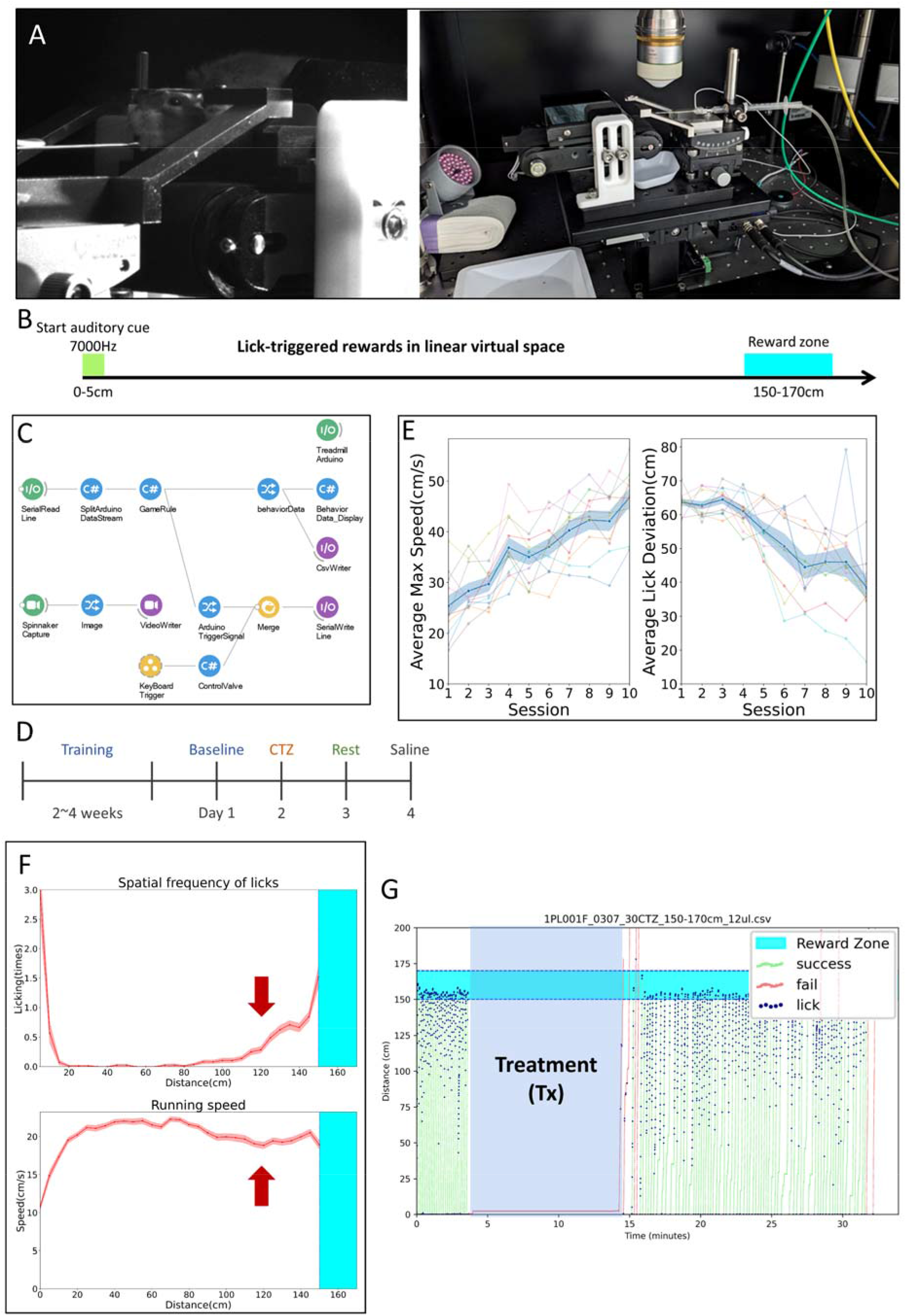
Self-motion-based path integration task design with nonspatial-related auditory cues allowed assaying dead reckoning for rewards using fine proprioceptive feedbacks. **(A)** VR system. Left, head-fixed mouse running on treadmill. Right, Overview of the apparatus that directly interacts with the animal. **(B)** Self-motion-based path integration task with nonspatial auditory cues allowed assaying dead reckoning for rewards using fine proprioceptive feedback. **(C)** The Bonsai workflow used for the auditory virtual environments treadmill task. **(D)** Experimental schedule. **(E)** Daily profile of average max speed and lick deviation over training. One session conducted per day. The blue line represents the population average with SEM, while the semi-transparent colored lines show individual data. The average maximum speed was calculated as the mean of the maximum speed for each trial in a session. Lick deviation was defined as the average distance between the reward zone floor (150 cm) and the center of mass (COM) of the licking positions in each trial. n = 10. **(F)** Representative example of lick and speed vs. virtual position from a well-trained mouse. This represents data from a single animal. The red line indicates the average of each trial within a session, with SEM shown as error area. The cyan area represents the reward zone. **(G)** Example of behavioral trajectory and lick patterns during a test session with CTZ injected to proximal limbs as the treatment. The semi-transparent blue area indicates the time during which treatment was conducted (no injection, saline, or CTZ). The green line represents the movement trajectory of the mouse in trials where a reward was obtained, while the red line represents trials without getting reward. Dark blue dots indicate the positions where the mouse licked.

### Memory-directed behavior for locating rewards in virtual space was affected by LMO3 perturbation

We wanted to test the function of proprioception in the context of relatively challenging goal-dependent behavior using the locally activated LMO3 system. The VR-like task was designed in such a way to solely rely on self-motions, functionally consistent with a process known as dead reckoning^38^. This setup included a virtual linear track and a spatially localized reward (**Fig. 7B**). Water-restricted mice were head-restrained, trained to navigate the environment with essentially no external landmark information with respect to the virtual space and no acceleration cues (because of head restraint), and lick within a narrowly defined RZ for obtaining a drop of sucrose solution.

With operant-conditioning design for spatially dependent reward-seeking, mastery of the task, evidenced by smooth running and anticipatory licking, typically required 2–4 weeks (**Fig. 7D**). During the training, animals demonstrated an increasing familiarity with the treadmill, reflected by a progressive increase in running speed over days, reduced uninstructed pausing, as well as concentration of the positions of licking closer to the RZ (**Fig. 7E**). Trained mice exhibited a consistent running speed drop and anticipatory licking as they approached the RZ (**Fig. 7F, G**), suggesting a capacity to rely on self-motion-based information for dead-reckoning. In principle, animals can solve the task by at least two kinds of strategies, namely spatial and temporal. The former relies on integration of the biomechanics during (virtual-spatially defined) self-motion activities, while the latter utilizes the stereotypic duration needed for each lap (trial) as long as over-trained mice achieve highly repetitive patterns of running across trials. Our analysis suggested that mice naturally develop a strategy using mixed information of those two for their path integration, which is, however, not a main emphasis here (and thus the data are not shown).

To analyze the effects of intramuscular LMO3 perturbation, we used two types of controls: pharmacological and genetic. Three metrics were used to assess the behavior, with delta (Δ) indicating the difference between pre-injection and post-injection measure:

1. **Ä Center of mass (COM) of lick positions from the start of the RZ (the 150^th^ cm)**, presumably reflecting the distance between the reward location anticipated by the animal and the true reward location
2. ***Within*-trial Δ Standard deviation (std) of lick positions**, presumably reflecting a (empirical) certainty of spatial inferences at the single-trial timescale
3. **Cross-trial Δ std of the mean COM of lick positions**, presumably reflecting a (empirical) certainty of spatial inferences at a cross-trial (20 trials) timescale

To measure differences during stable behavior, the first 10 trials immediately after the animal was returned to the treadmill following the treatment (Tx, body-restrained intramuscular CTZ injections) were excluded for analyses. Trials 11 to 30 (Pre-Tx) served as the baseline for comparison made with Trials 41 to 60 (Post-Tx) (**Fig. 8**). Under a relatively low-dose CTZ (0.1 mg/kg body weight) injected into the distal limb muscles, no significant differences were observed between pharmacological treatments **(Fig. S3)**. Similarly, under a relatively medium-dose CTZ (2.5 mg/kg body weight) injected into the distal limb muscles, no significant differences were detected between pharmacological and genetic treatments **(Fig. S4)**. In parallel, because of an earlier concern that injections into limb muscles affected the post-injection motivation due to discomfort, *i.p.* injections were also tested; no significant differences were found **(Fig. S5)**.

**Figure 8.**
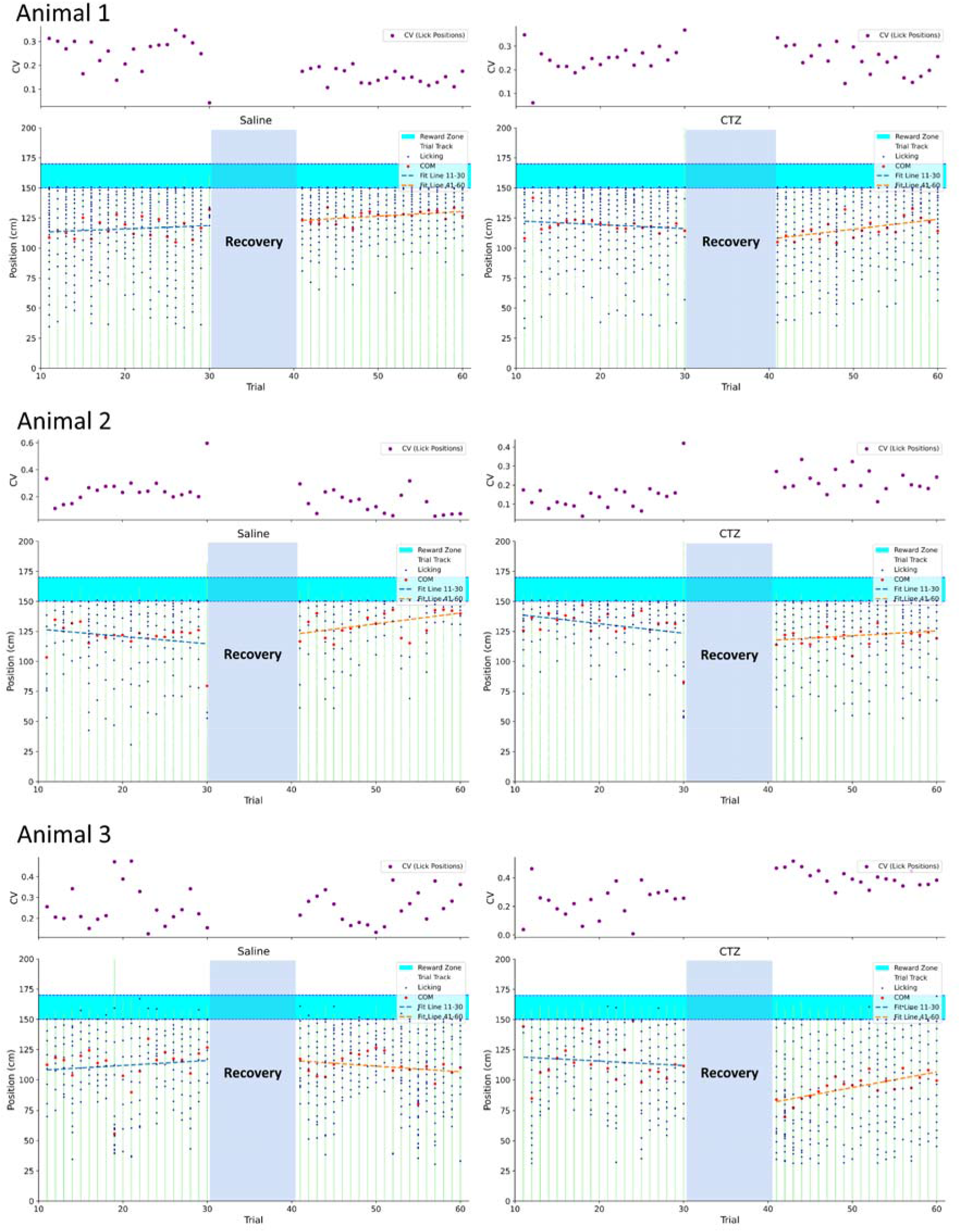
Example Spatial Lick-Pattern Dynamics from the Dead Reckoning Task. To compare behavioral differences with stable behavior, the first ten trials at the start of the task and immediately after the animal returned to the treadmill following treatment (Tx) were excluded (data not shown). Trials 11 to 30 (Pre-Tx) served as the baseline for comparison with trials 41 to 60 (Post-Tx). The left panel represents an individual animal‘s saline session, while the right panel shows the CTZ session. The green line illustrates the mouse’s position trajectory. Blue dots indicate the positions of lick events, and red dots represent the center of mass (COM) of lick positions. The cyan-colored block marks the reward zone. The blue and orange dashed lines represent the fitting of the lick COM distributions for pre-Tx and post-Tx, respectively. The coefficient of variation (CV) for lick positions in each trial is plotted on a separate axis above each subplot. Each purple dot represents the CV of a single trial.

Interestingly, when the high dose of CTZ (8.2 mg/kg body weight) was injected into the proximal limb muscles, PV-Cre;LMO3^f/f^ mice exhibited a significant increase in the distance between the lick-position center-of-mass (COM) and the RZ following CTZ, relative to their own saline session (**Fig. 9**; see also **Fig. 8**; paired mean difference = +9.2 cm; 95% CI [4.0, 13.7]; permutation *p* = 0.007; *n* = 12 mice). The within-trial and across-trial variability of lick-position COM showed the same direction of changes, but did not reach statistical significance (paired mean difference = +2.8 cm, *p* = 0.28; and +3.2 cm, *p* = 0.32, respectively). Notably, that none of these measures any nearly approached significance in the pharmacological control (saline) or in the genetic control animals, given the same CTZ injections, indicates a selectivity of the local manipulation: with the effect apparent only to a specific behavioral metric in a specific combination of transgenics and pharmacology, consistent with the notion of local luminopsin-based proprioceptive perturbation.

**Figure 9.**
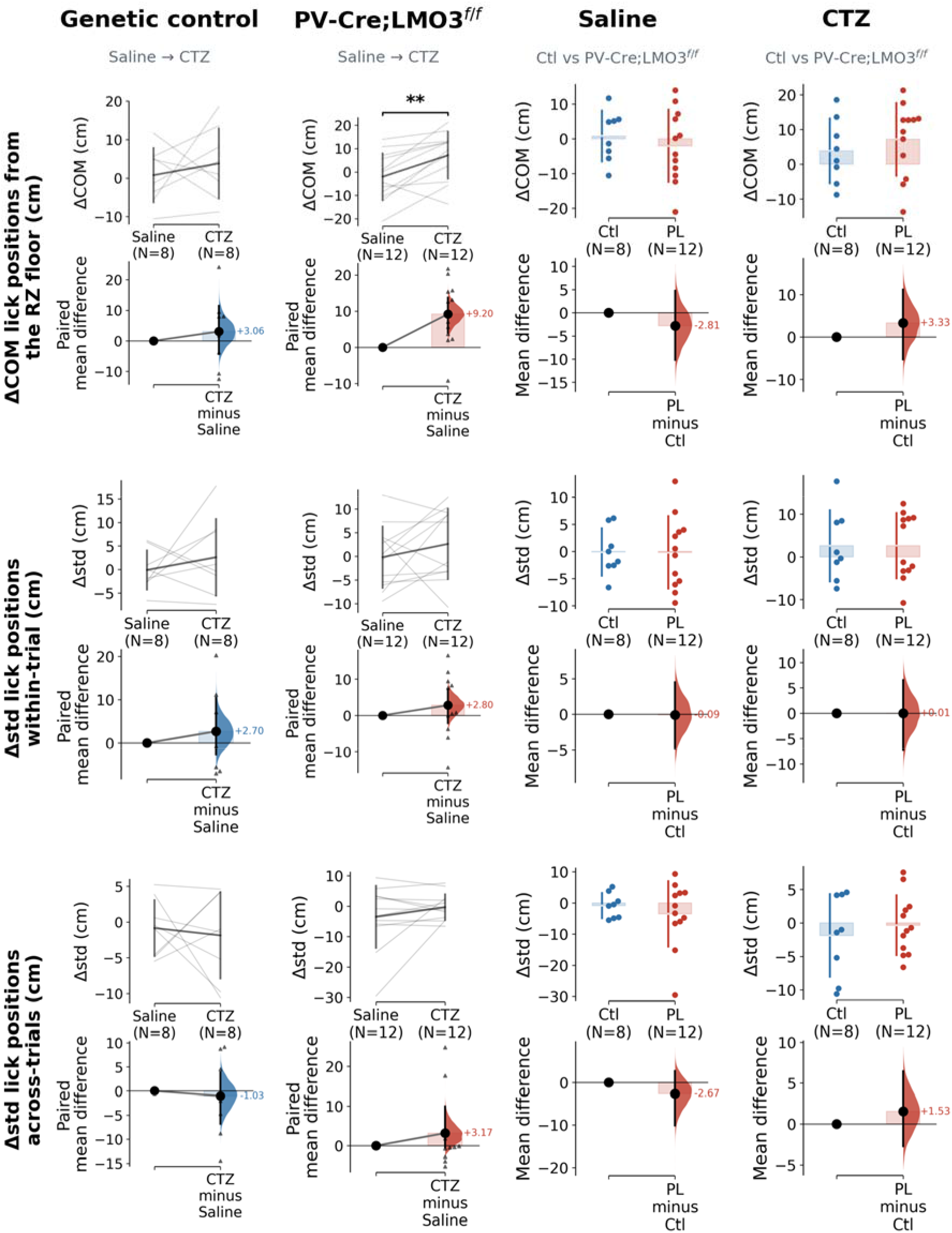
Proximal-limb injection of a high dose of CTZ (8.2 mg/kg body weight) produced a significant increase in anticipatory lick-position error in PV-Cre;LMO3^f/f^ mice. The Cumming estimation plot in this figure is similar in format to Figure 5 but includes paired lines and omits the vertical ends of the gapped lines representing the mean (gap) and ± standard deviation. ΔCOM of lick positions from the reward zone(RZ) “floor” was defined as the difference between pre-treatment (pre-Tx) and post-treatment (post-Tx) trials in the average distance between the reward zone floor (150 cm) and the center of mass (COM) of the licking positions in each trial. Δstd within-trial was defined as the difference (post-Tx minus pre-Tx) in the average standard deviation of ΔCOM of lick from RZ within a single trial. Δstd cross-trials was defined as the difference (post-Tx minus pre-Tx) in the average standard deviation of ΔCOM of lick from RZ across trials, with 20 trials analyzed separately for post-Tx and pre-Tx. Ctl n = 8 mice (genetic controls); PV-Cre;LMO3^f/f^ (PL) n = 12 mice. Within-genotype Saline-vs-CTZ comparisons used a paired permutation test; between-genotype comparisons used an unpaired permutation test (5,000 resamples, DABEST). ** *p* < 0.01. Blue, genetic control; red, PL.

We further examined the effects of CTZ dosage on licking behavior. The change in the COM of lick positions (ΔCOM) exhibited a trend of dose dependence, with higher doses tending to induce a larger spatial shift, but the difference across the four dose × injection site conditions did not reach statistical significance (p = 0. 739, **Fig. S6A**). In contrast, the metrics of licking variability—both within-trial (p = 0. 933, **Fig. S6B**) and between-trial (p = 0. 891, **Fig. S6C**)—showed no clear dose-dependent effects, nor significant differences between injection sites. Notably, the magnitude of the treatment effect did not correlate with each animal’s baseline licking variability, suggesting that the observed change in licking metric was less explained by known intrinsic linking characteristic associated with individual animals (**Fig. S7**).

From our empirical observation, the gentle awake-constraint procedure still produced certain levels of interference on mice’s post-injection mobility. We thus consider if the running dynamics has anything to say about the spatial positions of licking, or can even reflect, in any case, proprioceptive luminopsin perturbation on its own. Under CTZ, however, post-injection running speed was statistically indistinguishable between PV-Cre;LMO3^f/f^ and control mice (mean speed Δ = +0.14 cm/s, p = 0.92; peak speed Δ = +0.89 cm/s, p = 0.65; permutation test). On the other hand, the fraction of immobile time (speed < 4 cm/s, measured within the 20-min post-injection window) did not differ between control and PV-Cre;LMO3 ^f/f^ under CTZ injections, either (Δ = −0.05, p = 0.41; permutation test). Therefore, the genotype- and ligand-specific anticipatory-lick change is less likely to be attributed to any change in locomotor speed or moving time.

Taken together, across our extensive screening across combinations of tasks, transgenics, pharmacological (ligand) conditions and dosages, these results suggest that the LMO3 system may interfere with the animals’ dead reckoning ability by potentially increasing errors in spatial inference, an effect more prominent than the measured inference certainty proxies possibly due to securing of the rewards still with the dispersed lick patterns as well as (potentially) the trial time still a reliable contingency signal useful for the task. We used the established tissue-specific luminopsin transgenetic mouse line, not only as a proof-of-concept of mild perturbation tactics for navigation-relevant proprioception, but also as a first temporally controllable, selective sensory manipulation for path-integration processes in behaving mammals, to our understanding.

## Discussion

### Critical scientific need for animal models allowing unambiguous and transient manipulation of proprioception

Proprioception plays a pivotal role in establishing the egocentric, and its conversion to allocentric, frame of reference in spatial cognition^39^. Body perception is seen indispensable for memory and intelligent embodied control in space^40^. Understanding how neural circuits enable spatial navigation requires uncovering the mechanisms by which proprioception facilitates encoding of bodily states and egocentric representations^5^. Despite the importance, progress in proprioception research has been hindered by the absence of animal models and paradigms that allow selective and reversible manipulation of proprioceptive pathways. Current approaches, highlighted in recent studies^15,20,41^, often rely on indirect and complex methodologies or theoretical modeling, with certain limitations on clear biological causality and mechanistic clarity. Thus, the development of experimental approaches capable of directly, selectively and reversibly manipulating proprioceptive functions is imperative.

LMOs are bioluminescent opto-chemogenetics tools that integrate the strengths of optogenetics and chemogenetics. Optogenetics offers exceptional spatiotemporal precision but is limited by a need for invasive light delivery^42–44^, while chemogenetics enables neural modulation without optical implants but has biochemically delimited slow onset and prolonged mode of action, due to G-Protein-Coupled Receptor-based signaling^45,46^. In contrast, we chose luminopsins because they provide optogenetics-like precision via bioluminescence, eliminating external light sources, and have better temporal control through *in-vivo* pharmacokinetics of luciferase ligands and actions of ion channels^26^. In the future, even different modalities of control (opto or chemo) could be applied in the same animals, providing results from different manipulations with a same, comparable baseline.

### Route of administration for the luciferase ligand (CTZ) and the pharmacodynamics

A critical issue of this new peripheral modality-selective manipulation is the duration and strength of the effects. Research using LMOs to modulate neural activity during animal behavior is still limited. Previous studies, including those on tool development and neural regeneration, have employed various administration routes (such as intracranial, intravenous(IV), intraperitoneal (IP), and intranasal injections (IN)) and dosages ^26,31–34,47–49^. Intravenous CTZ in rodents induces rapid effects, with spike rates increased within seconds, peaking at 20–30 seconds, and decaying over minutes^26^. Intranasal delivery offers quick brain targeting within minutes^31^. Intraperitoneal delivery, though slower, is preferred for repeated applications, with effects peaking within 5 to 40 minutes and lasting up to an hour. Bioluminescence intensity and duration are dose-dependent, with higher doses producing stronger and longer-lasting effects. For instance, a 20 mg/kg dose yielded approximately 4 times the intensity and duration of a 5 mg/kg dose^31,33,34^.

In this study, we employed local intramuscular (IM) injections to selectively modulate the peripheral source of proprioceptive signals in limbs. The relatively shorter duration of observed effects (as in the Grid Walk), as well as sensitivity to only a subset of behavioral tasks, is consistent with a localized, mild manipulation without wider-spread neural effects, such as gross motor coordination, postural regulation and basal motion kinematics, effort-based motivation, and longer-term memory of task rules. Not as the effects seen in the mice with genetic knockout of acid-sensing ion channel subunit type 3 (ASIC3^27^)—which showed more missed stepping (foot faults) in Balance Beam—local LMO3 activation produced no pronounced effects in the same task.

This discrepancy may be due to CTZ-dosage-, site-of-injection- or LMO-variant-dependent insufficient excitation of PV^+^ cells in theDRG,. Future studies could improve outcomes by using modified strategies or non-invasive opto-stimulation with more sensitive opsin variants. Nevertheless, a mild enough control is still of highest strategic priority: a real challenge for studying proprioceptional computation in spatial cognition is a relative isolation of tool effects from acting on apparent motor-control functions.

### Do different behavioral tasks involve proprioceptive functions in different ways?

No behavioral effect of LMO was observed in the Balance Beam task, and the Open Field showed little specific enough change (except for increased left-turning, which, however, was not temporally specific to the inferred effective CTZ duration of ∼15 mins; **Figure S1F**) with general locomotor capacity, kinematics and explorative behaviors intact, whereas statistically significant differences were observed in the Grid Walking and Dead Reckoning tasks. One possible explanation is that different behavioral tasks involve proprioceptive functions in distinct ways. Proprioception is a system that integrates multiple components (including muscle tension, movement speed and joint angles^17,50,51^) underlying multidimensional subtle motor control, body image and schema^40^, thus unlikely to contribute in various behavioral contexts the same way.

In this context, the Balance Beam and Grid Walking tasks examined an animal’s ability to sense and fine-tune its body movements under different conditions. In our experiment, CTZ was injected into all four limbs (which is different from Lin et al.^27^). In Balance Beam, the mice might be able to adapt to the relatively symmetric impact on all limbs. In contrast, Grid Walking, in darkness, relied less on whole-body balance and was more specifically about the individual limb’s positioning and fine maneuvers. This difference might explain the differential sensitivities of task performance to LMO3 activation. An experiment with unilateral CTZ injection can serve to test this hypothesis. Overall, proprioceptive signal types and precision required for these tasks likely differ fundamentally.

### Dead Reckoning task could provide a readout for neuro-mechanics support of behavioral licking decision in virtual space

The Dead Reckoning task potentially assessed whether the animal can integrate its past movements to make decision on licking. Thus, the task may reveal significant differences in specific, spatial behavioral metrics that were not accounted for by the subtle differences seen in simpler tasks, necessitating a closer deliberation of the interpretations on each measure and the observed statistical significance.

ΔCOM of lick from the RZ was defined as the difference between pre-Tx and post-Tx trials in the average distance between the RZ floor (150 cm) and the COM of licking positions in each trial. This measure reflects the animal’s ability to adapt its reward location prediction based on integrated sensory and motor feedback across two treatments. A larger ΔCOM indicates greater deviation from the expected reward position, suggesting impaired spatial integration.

Δstd within trial was defined as the difference (post-Tx minus pre-Tx) in the average standard deviation of ΔCOM of lick from RZ within a single trial. This measure captures changes in the variability of licking behavior within a trial between the two treatments. Increased variability within a trial may indicate inconsistent integration of proprioceptive and external sensory cues.

Finally, Δstd between trials was defined as the difference (post-Tx minus pre-Tx) in the average standard deviation of ΔCOM of lick from RZ across trials within a session. This measure reflects changes in trial-to-trial behavioral consistency between the two treatments. A higher Δstd between trials suggests instability in forming a reliable internal spatial representation across repeated attempts.

The observed statistical significance in the spatial dead reckoning task, in contrast to tasks like Grid Walking or Balance Beam, may be attributed to the unique demands of integrating proprioceptive signals with decision-making processes. While tasks such as Grid Walking and Balance Beam primarily assess real-time proprioceptive feedback and fine motor adjustments, the dead reckoning task requires animals to encode, store, and retrieve spatial and motion information over time to guide behavior. This additional computational complexity likely amplifies subtle deficits in proprioceptive signal integration, making them more detectable in our experimental design.

### Limitations of the study and future work

This study offers a highly promising animal behavior system that allows precise control over the information available to animals, as well as preliminary insights into the role of proprioception in spatial navigation. However, several limitations must be addressed to enhance the validity and generalizability of the conclusions.

A key limitation is the exclusive reliance on LMO3. While LMO3 effectively disrupts proprioceptive signaling, it may not achieve sufficient activation of target neurons in all contexts. Future research could explore enhanced LMO variants with improved efficacy and specificity to enable more robust modulation of proprioceptive pathways^49,52^. Additionally, the CTZ dosage and delivery method require refinement. Higher doses or injection strategies targeting areas with greater proprioceptor density could potentially amplify the effects on proprioceptive signaling. Alternatively, emerging neuromodulation tools offer potential alternatives. For instance, ChRmine, a novel red-shifted channelrhodopsin, enables neuronal activation without the need for invasive light sources. This approach has been successfully validated in deep brain regions, such as the ventral tegmental area, located 4.5 mm beneath the skull surface^53,54^. ChRmine could potentially replace LMOs, providing a less invasive method (eliminating the need for injections) and higher temporal precision (bypassing pharmacokinetic limitations) for modulating proprioceptive neurons at muscle spindles. Additionally, Myomatrix arrays^55^ or magnetogenetics could also serve as potential alternative solutions.

To independently validate the findings, future studies will employ complementary methods, such as genetic manipulation of other proprioceptive pathways (e.g., ASIC channels)^27^. Moreover, advanced imaging techniques, such as two-photon microscopy, could directly visualize neuronal activity changes resulting from LMO modulation, providing more detailed mechanistic insights.

By addressing these limitations and implementing the suggested refinements, future studies can build upon the foundation of this work to further elucidate proprioception’s contributions to spatial cognition.

---

**Supplementary Figure 1.**
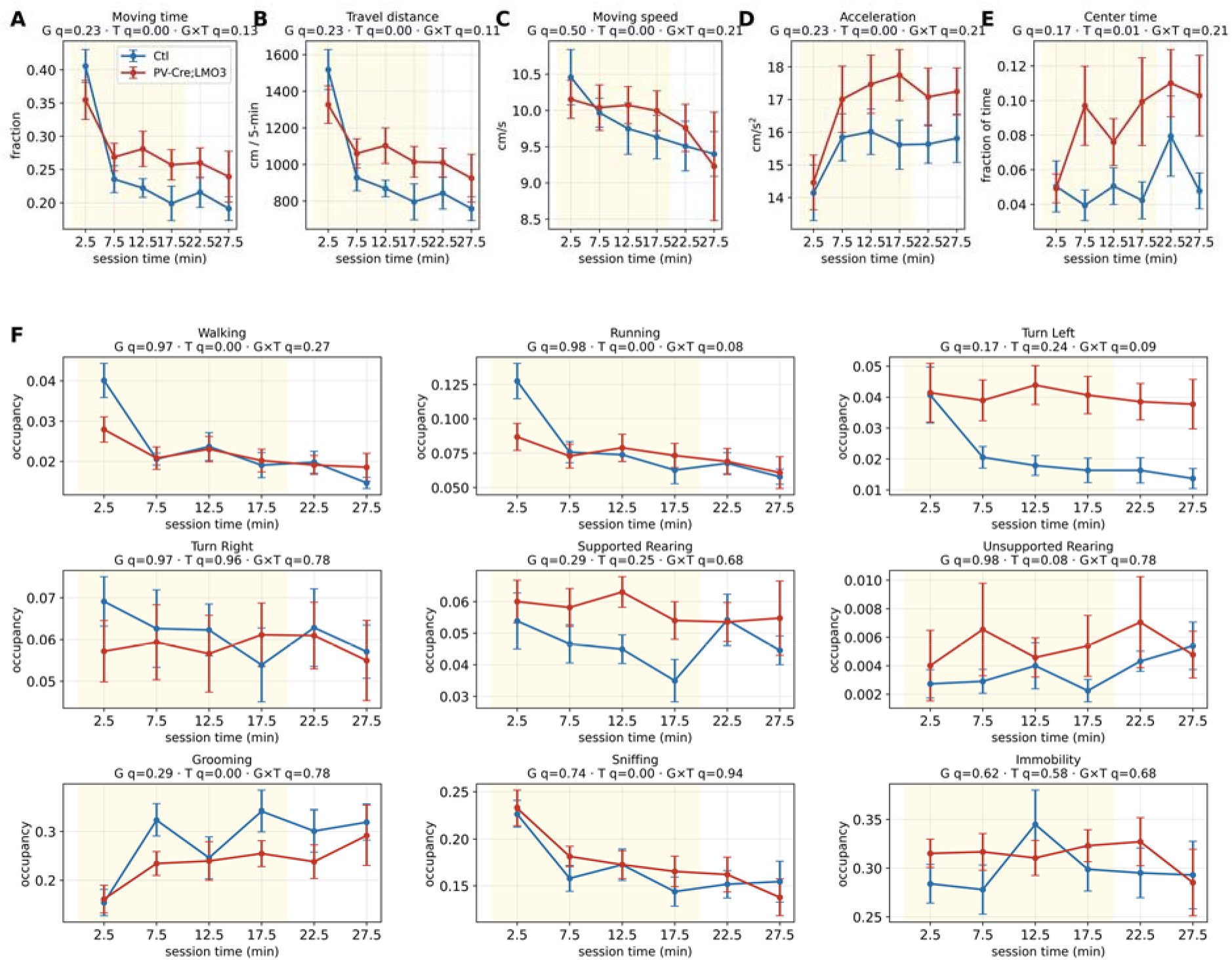
Lack of significant effects of luminopsin activation on behaviors in Open Field with temporal specificity over the full 30-min session of recording Complementary to Figure 2, with the same analysis and visualization as Figure 5D. Ctl, blue, *n* = 11; PV-Cre;LMO3^f/f^, red, *n* = 14. Shaded pink region, estimated CTZ effective time window based on IVIS bioluminescent measurement. Due to post-handling neuro-behavioral instability, statistical analysis was performed only over 5-30 mins. (A–E) Similar to Figure 2A-E, but over post-injection time. (F) Similar to Figure 2J, but over post-injection time. Aligned-rank-transform (ART) two-way repeated-measures ANOVA; subtitles showing Benjamini–Hochberg FDR-corrected *q* value for the genotype (G), (T) and genotype × time interaction (G × T) effect. No significance was found in genotype × time interaction a test for genotype-specific temporal changes, suggesting no temporally-specific LMO3 activation on Open-Field behaviors was observed. \**q* < 0.05. At the first 5-min time bin (0–5 min), group differences in per-class occupancy were tested by unpaired permutation of the mean difference (5,000 resamples) with Benjamini–Hochberg correction across the nine classes. Walking (Ctl 0.040 vs PV-Cre;LMO3 0.028): Δ = −0.012, p = 0.030, q = 0.134. Running (0.128 vs 0.087): Δ = −0.041, p = 0.020, q = 0.134. Both were nominally lower in PV-Cre;LMO3 at 0–5 min (uncorrected p < 0.05) but did not survive FDR correction (q = 0.13); no class reached FDR significance at this time bin.

**Supplementary Figure 2.**
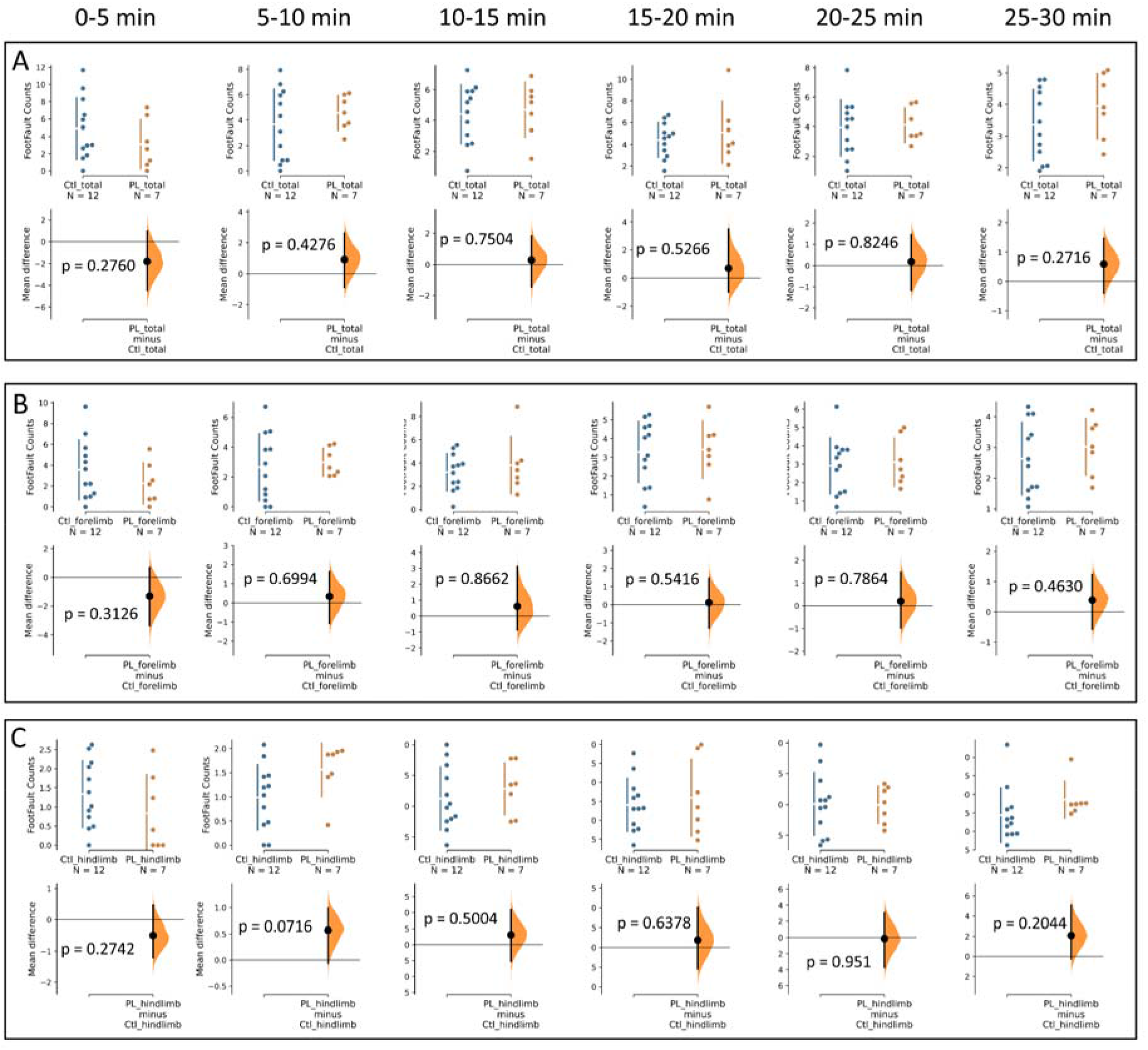
Visualization of individual time bin data and post-hoc analysis for the grid walking test. The Cumming estimation plot in this figure is similar in format to Figure 5. Ctl (LMO3^f/f^) n = 12 mice, PV-Cre;LMO3^f/f^ n = 7 mice; Unpaired permutation test; no significant difference; n.s.. **(A)** Cumming estimation plots comparing all foot faults across each time bin. **(B)** Cumming estimation plots comparing forelimb foot faults across each time bin. **(C)** Cumming estimation plots comparing hindlimb foot faults across each time bin.

**Supplementary Figure 3.**
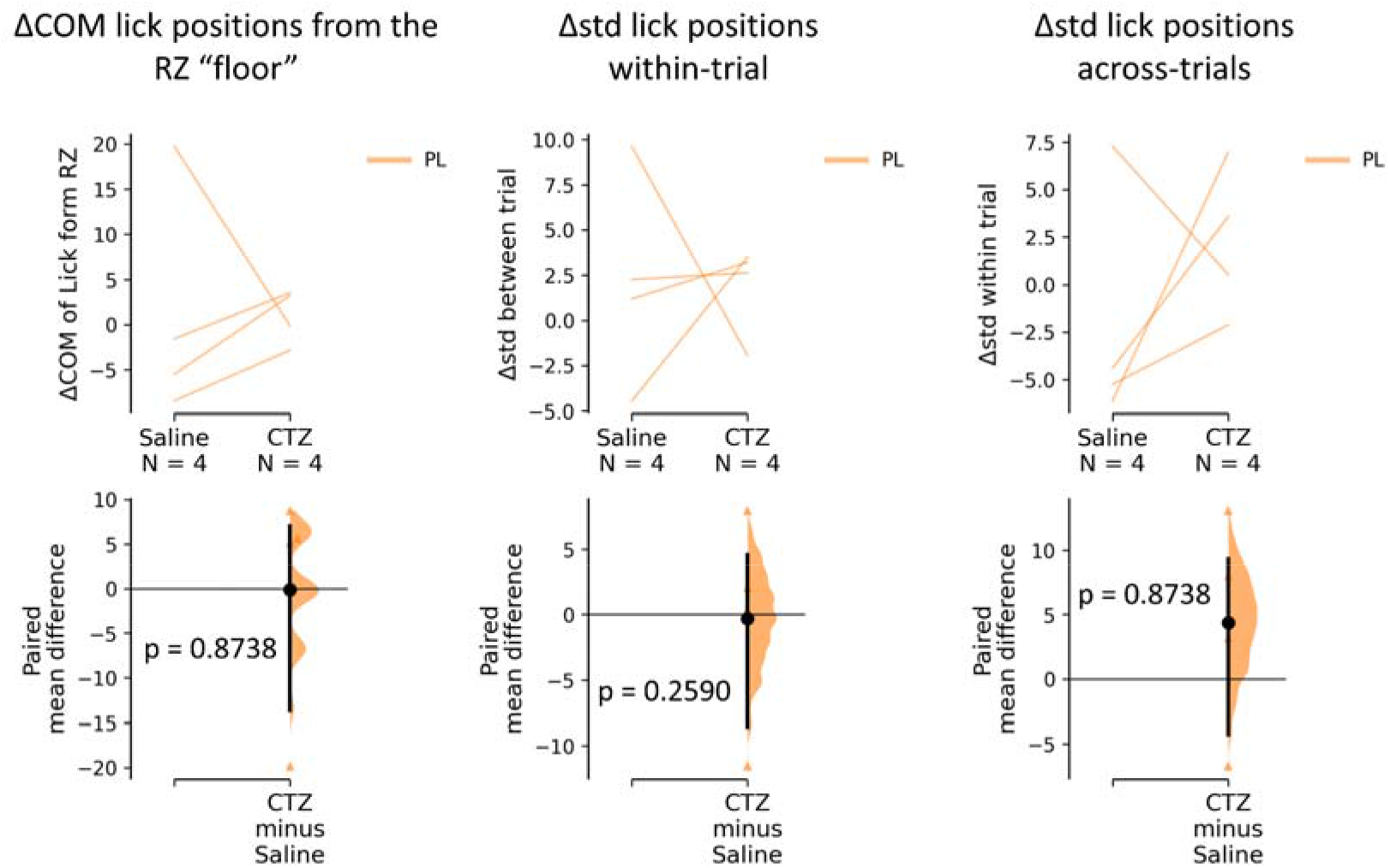
Mice with distal muscle injections of CTZ at a relatively low dose (0.1 mg/kg body weight) showed no significant difference in anticipatory lick position or variance The Cumming estimation plot in this figure is similar in format to Figure 5&9. The behavioral metrics in this figure are the same as those presented in Figure 9. This experiment used the treatment as the control, and all mice were of the PV-Cre;LMO3^f/f^ (PL) genotype. PV-Cre;LMO3^f/f^ (PL) n = 4 mice; Unpaired or paired permutation test; no significant difference; n.s.

**Supplementary Figure 4.**
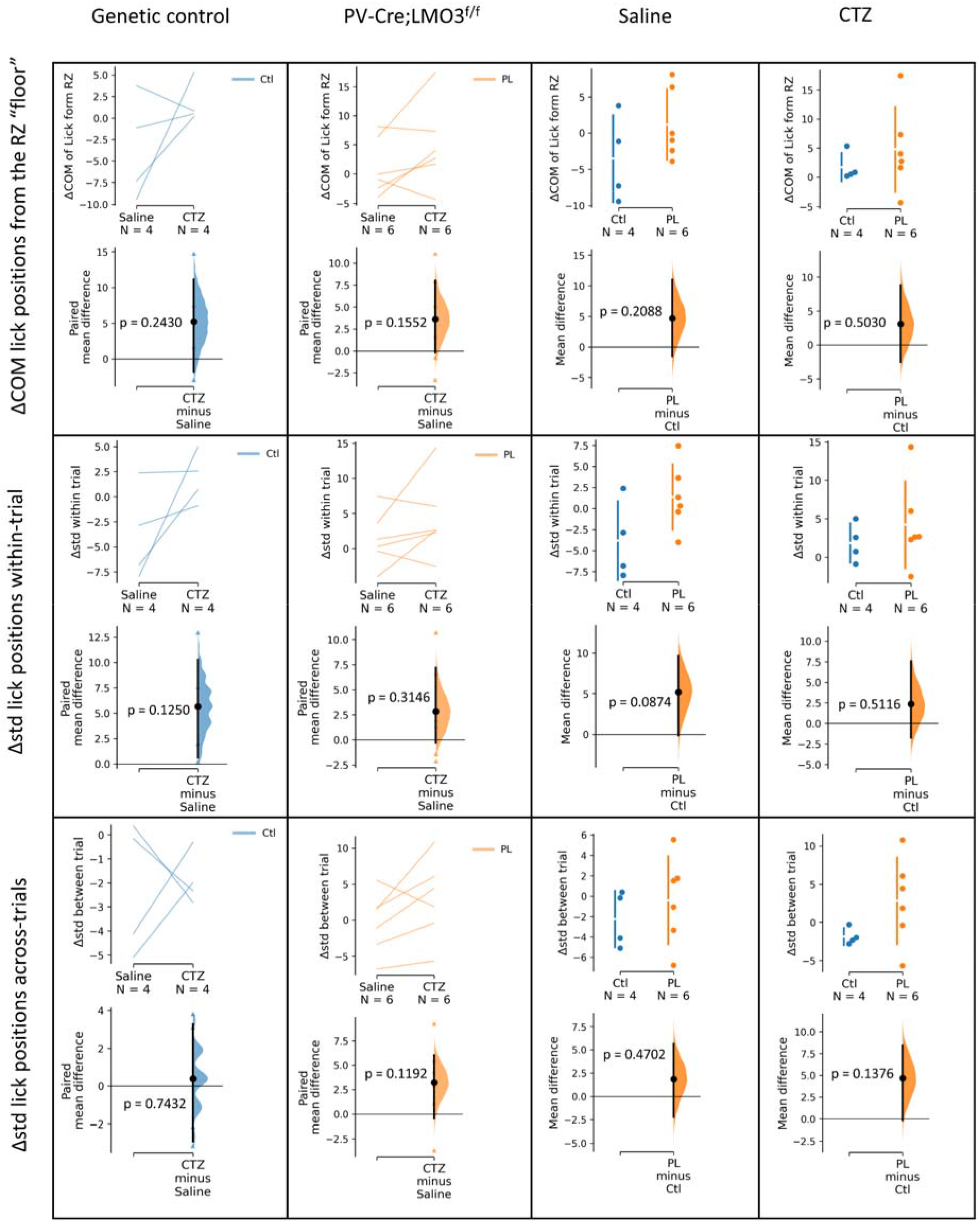
Mice with distal muscle injections of CTZ at a relatively medium dose (2.5 mg/kg body weight) showed no significant difference in anticipatory lick position or variance The Cumming estimation plot in this figure is similar in format to Figure 5&9. The behavioral metrics in this figure are the same as those presented in Figure 9. Here were comparisons of two different types of pharmacology and genotype control. Ctl (LMO3^f/f^) n = 4 mice, PV-Cre;LMO3^f/f^ (PL) n = 6 mice; Unpaired or paired permutation test; no significant difference; n.s.

**Supplementary Figure 5.**
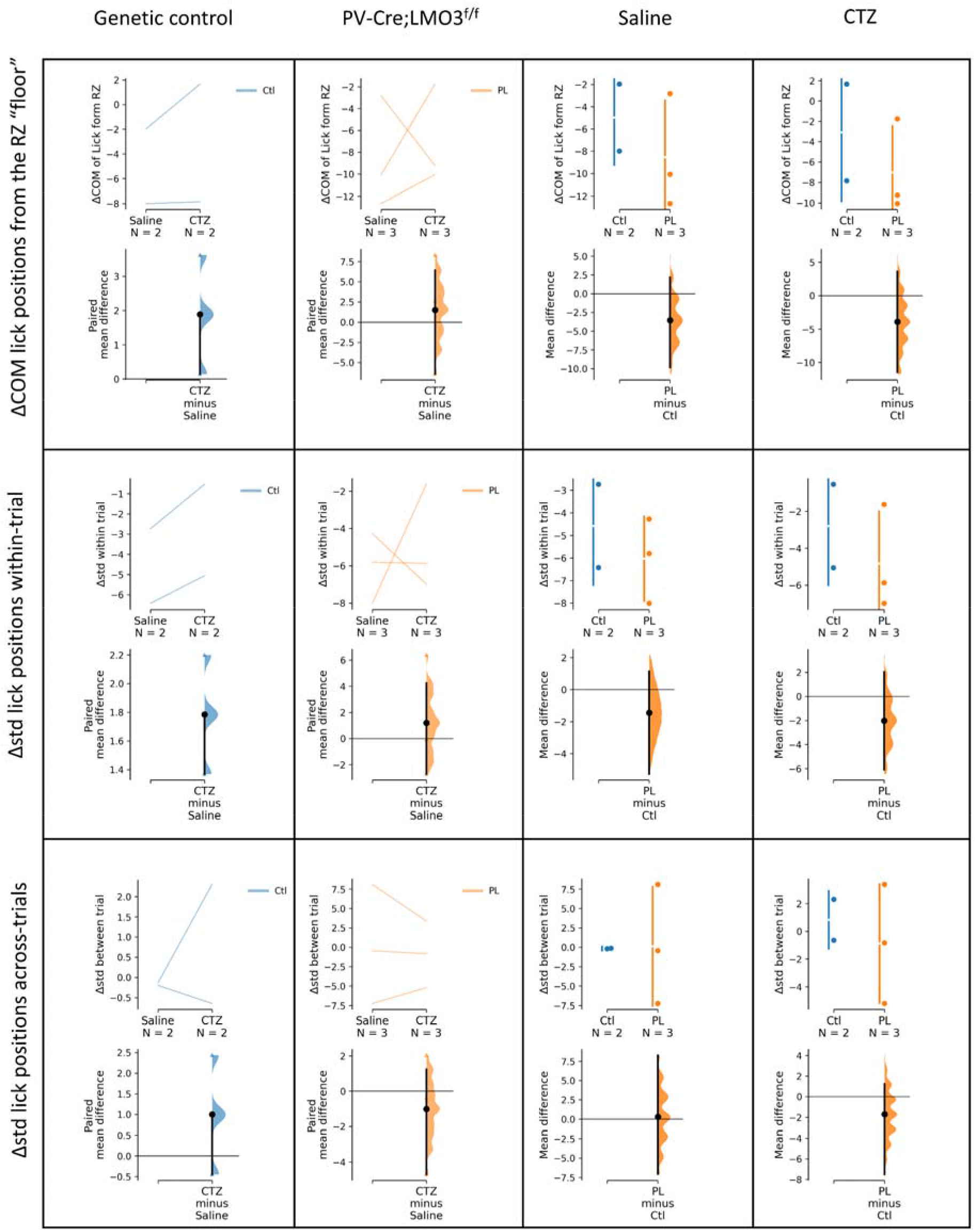
Mice with intraperitoneal injections of CTZ at a relatively high dose (20 mg/kg body weight) showed no difference in anticipatory lick position or variance (preliminary tests) The Cumming estimation plot in this figure is similar in format to Figure 5&9. The behavioral metrics in this figure are the same as those presented in Figure 9. Ctl (LMO3^f/f^) n = 2 mice, PV-Cre;LMO3^f/f^ (PL) n = 3 mice. Because the sample size was very small, statistical tests were not performed.

**Supplementary Figure 6.**
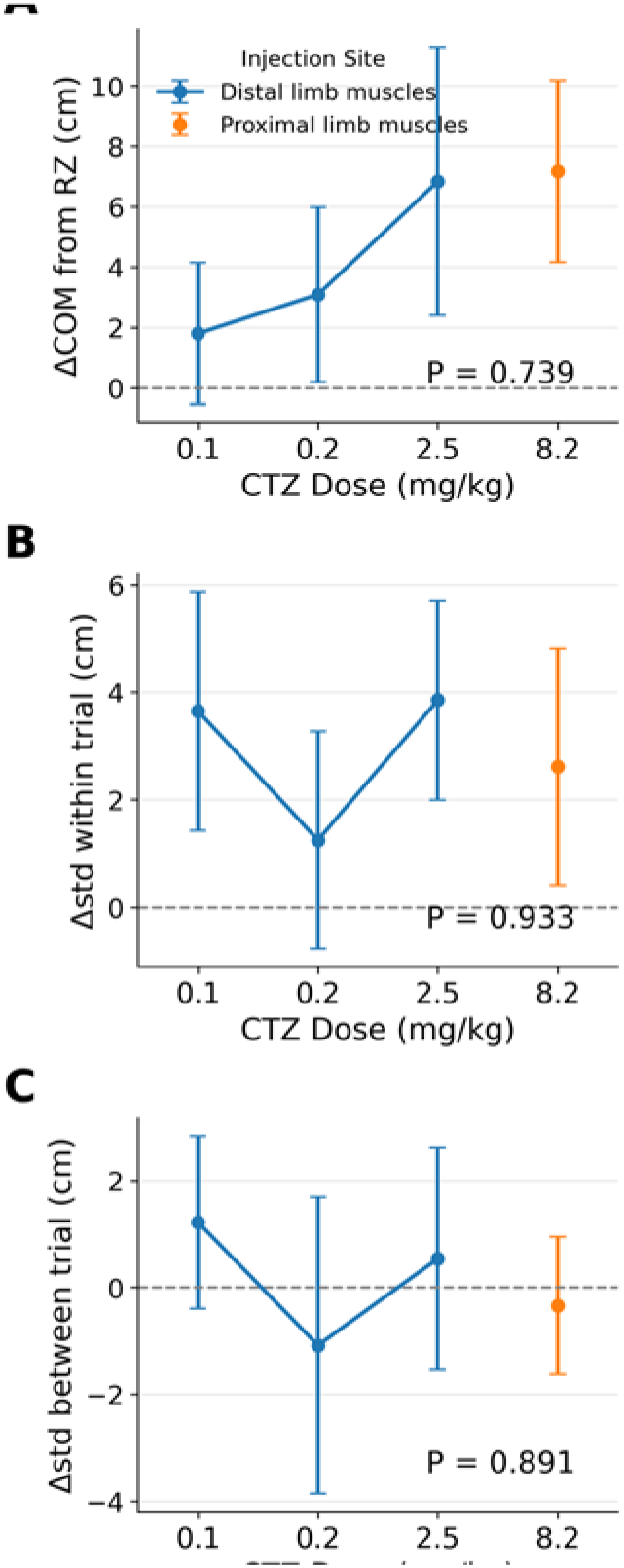
Effects of CTZ dose and injection site on licking behavior. The behavioral metrics in this figure are the same as those presented in Fig. 9, following CTZ injection into either distal (blue line) or proximal (orange dot) limb muscles. All mice were o f the PVCre;LMO3f/f (PL) genotype and only CTZ sessions are shown; each mouse contributes a single value per dose, computed over the same 20 preTx and 20 postTx trials used in Fig. 9. Distal muscle injection: 0.1 mg/kg body weight, n = 4 mice (Supplementary Figure 3); 0.2 mg/kg, n = 3 mice; 2.5 mg/kg, n = 6 mice (**Figure S4**). Proximal muscle injection: 8.2 mg/kg, n = 12 mice (Fig. 9). (**A**) ΔCOM of lick from RZ. (**B**) Δstd within-trial. (**C**) Δstd cross-trials. Across all three measures there was no significant difference across doses (one-way ANOVA: (A) *F*(3,21) = 0.42, *P* = 0.739; (B) *F*(3,21) = 0.14, *P* = 0.933; (C) *F*(3,21) = 0. 21, *P* = 0.891); n.s. All data are presented as mean ± SEM.

**Supplementary Figure 7.**
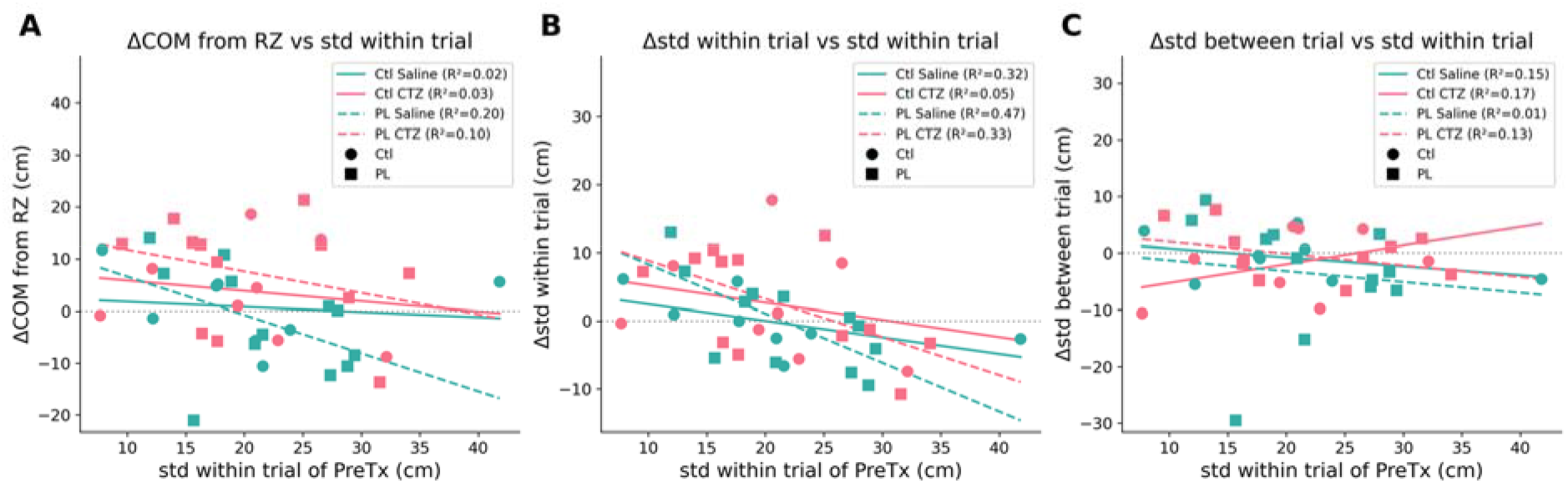
The effects of treatment did not correlate with baseline within-trial variability of licking. The behavioral metrics in this figure are the same as those presented in Fig. 9, and the animal s are the same cohort. For each of the four genotype × treatment combinations, the treatment effect (y axis, post-Tx minus pre-Tx) was plotted against that animal’s own pre-Tx within-trial standard deviation of lick position (x axis) and fitted by ordinary least squares; the R² of each fit is given in the legend. (**A**) ΔCOM of lick from RZ. (**B**) Δstd within-trial. (**C**) Δstd cross-trials. The combination in which proprioceptive feedback is expected to be perturbed — PL mice given CTZ — behaved no differently from the other three combinations: its regression slope carried the same sign as the control combinations in every panel (all four slopes negative in (**A**) and in (**B**); negative in (**C**), as in Ctl saline and PL saline), its slope magnitude fell wit hin the range spanned by them, and none of its correlations reached significance ((**A**) *r* = −0.3 1, *P* = 0.329; (**B**) *r* = −0.57, *P* = 0.051; (**C**) *r* = −0.36, *P* = 0.252; Pearson). Of the twelve fits, only one reached nominal significance — PL saline in (**B**) (*r* = −0.68, *P* = 0.014) — that is, in a vehicle condition rather than in the CTZactivated PL group. The direction and magnitude o f the treatment effect were therefore not accounted for by how variable an animal’s licking alr eady was before treatment, and this was equally true of the CTZ-activated PL mice.

## Key Resources Table

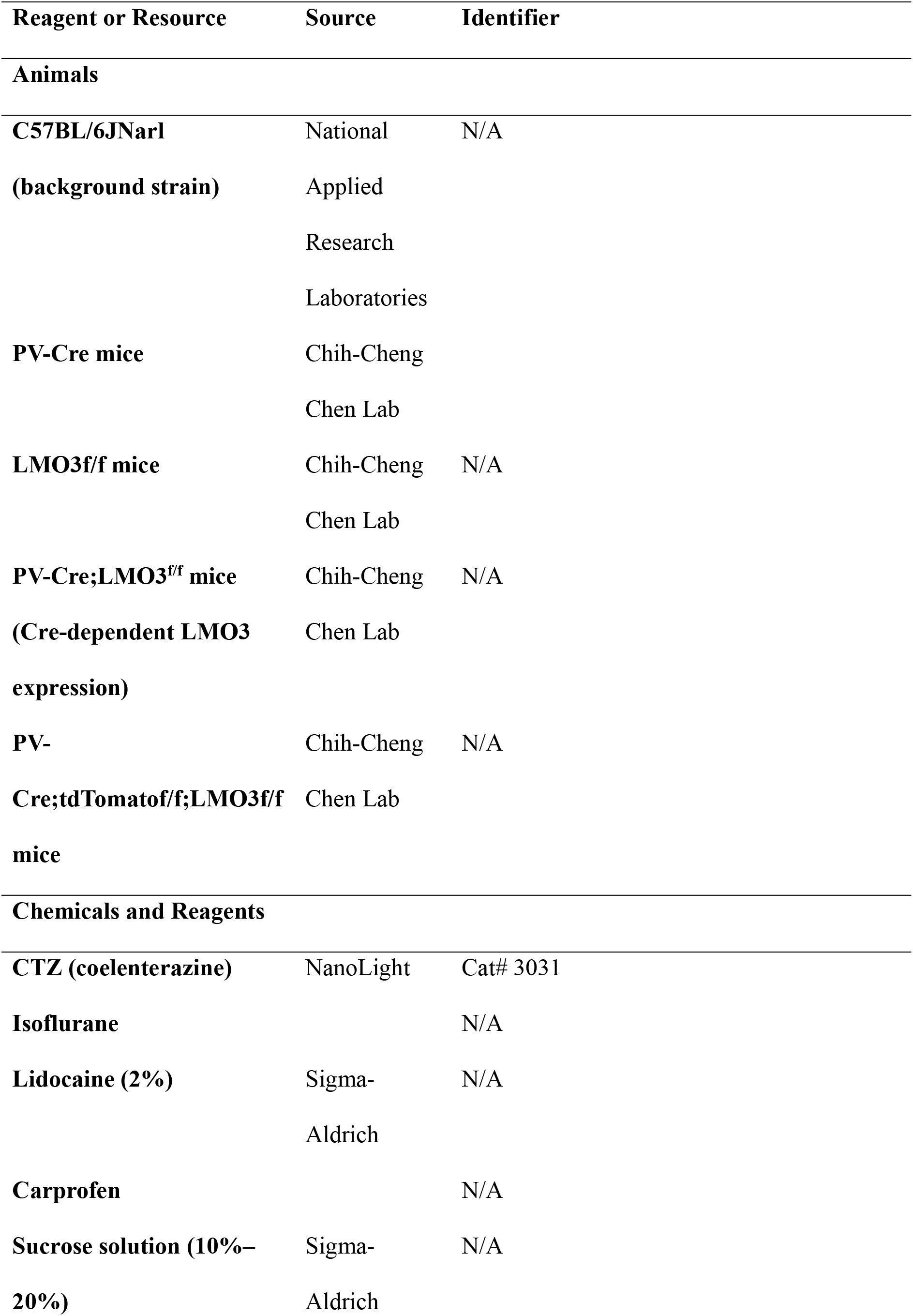
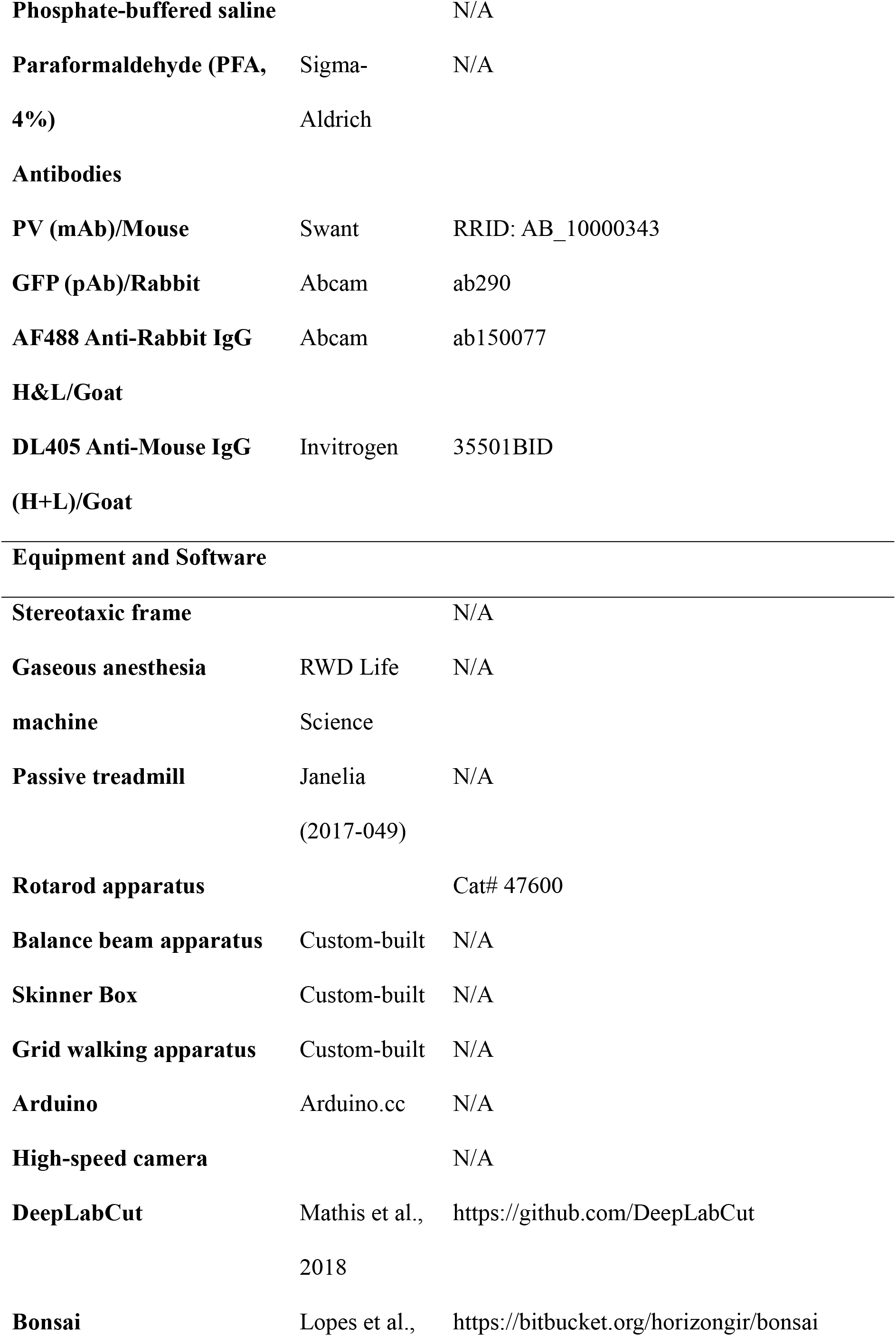
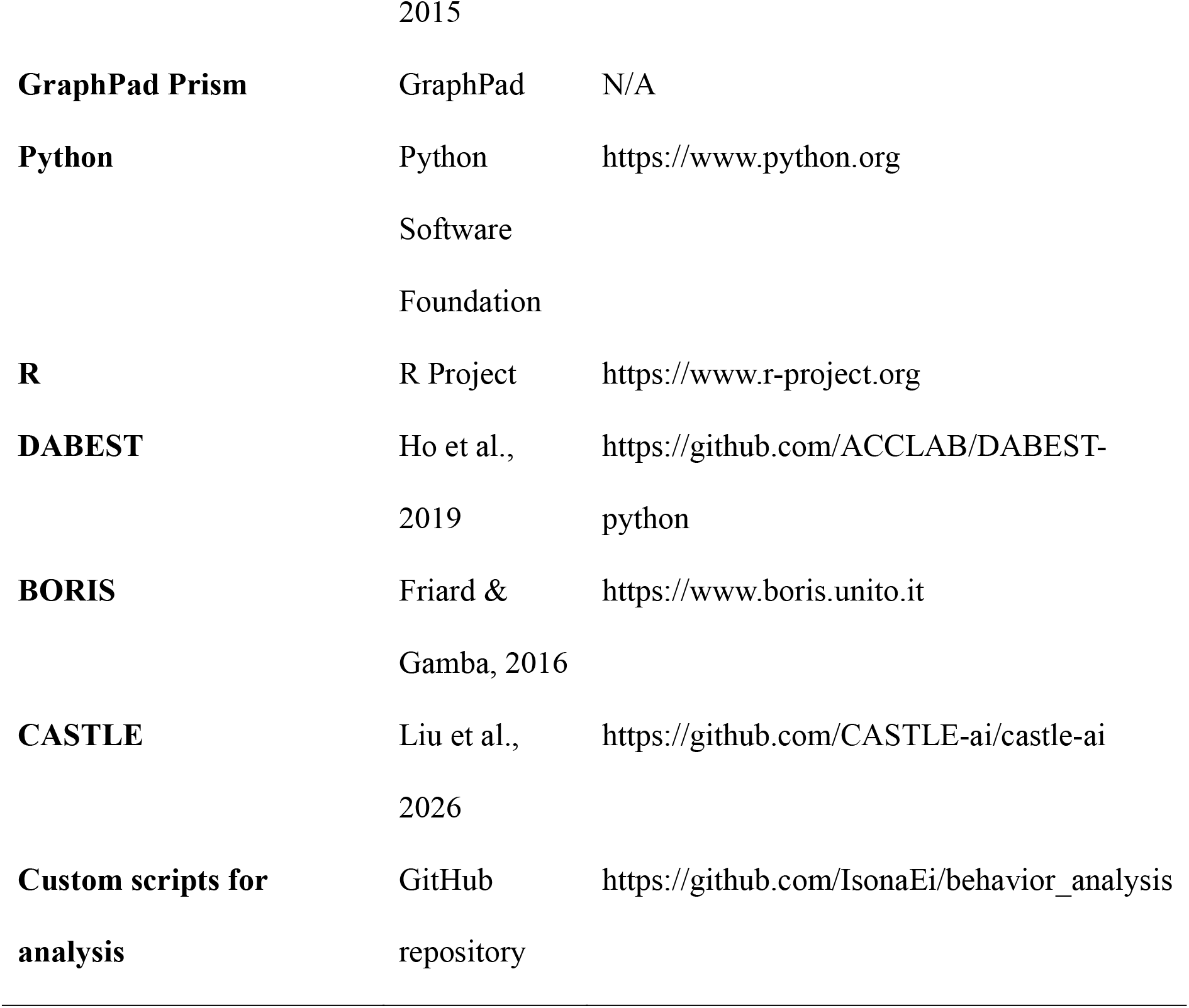

## RESOURCE AVAILABILITY

### Lead Contact

Further information and requests for resources, data, and reagents should be directed to and will be fulfilled by the lead contact, Ching-Lung Hsu.

### Materials Availability

This study did not generate new unique reagents.

All genetically modified mice used in this study are available upon request from the Chih-Cheng Chen Lab. The background strain C57BL/6JNarl is maintained by the National Applied Research Laboratories (NARLabs).

### Data and Code Availability

Data analysis scripts will be publicly available on GitHub at: https://github.com/Ching-Lung-Hsu-Lab/ProprioceptionPathIntegration

Python and R scripts used for statistical analysis and visualization are included in the repository, along with a README file providing step-by-step instructions.

Any additional information and datasets required to reanalyze the data reported in this paper is available from the lead contact upon request.

## EXPERIMENTAL MODEL AND SUBJECT DETAILS

All animals used in this study were genetically modified C57BL/6JNarl mice, a substrain established and maintained by the National Applied Research Laboratories (NARLabs, Taiwan). The following transgenic mouse lines were used: PV-Cre mice (Pvalbtm1(cre)Arbr; Jackson Laboratory, Stock #017320), which express Cre recombinase under the parvalbumin (PV) promoter; LMO3f/f mice (Tg(Hsyn-LMO3)tm1), generated by the Chih-Cheng Chen Lab, which carry a floxed allele of LMO3 for Cre-dependent expression. From these lines, PV-Cre;LMO3^f/f^ mice were generated to achieve Cre-dependent LMO3 expression in parvalbumin-expressing neurons. Additionally, a triple transgenic line, PV-Cre;tdTomatof/f;LMO3f/f mice, was generated by crossing PV-Cre, LMO3f/f, and Ai14 (tdTomato reporter) mice, enabling Cre-dependent expression of both LMO3 and tdTomato. The control groups consisted of PV-Cre mice only, which lacked LMO3 expression, and hSyn-LMO3f/f mice only, which did not undergo Cre recombination. All experimental mice were bred within the same colony to ensure consistent genetic backgrounds, and all procedures were approved by the Institutional Animal Care and Use Committee (IACUC) of Academia Sinica, adhering to both institutional and national guidelines for ethical animal research.

All animals were housed in the Animal Facility of the Institute of Biomedical Sciences, Academia Sinica in standard conditions with a temperature range of 20–23°C, relative humidity between 40%–70%, and a 12-hour light/dark cycle (lights on at 7 AM). Mice were housed in groups of up to five per cage and provided with environmental enrichment, including igloos, running wheels, wooden sticks, and cotton pads, to alleviate stress and encourage natural behaviors. Standard chow and water were available ad libitum, except for mice undergoing behavioral experiments requiring water restriction, where they were limited to 1 mL of water per day for at least three days prior to the start of training.

For surgical procedures, animals undergoing headplate implantation were first anesthetized with 3% isoflurane for induction, followed by maintenance at 1.5% isoflurane in pure oxygen. After securing the mice in a stereotaxic frame, the skull was exposed, and a stainless steel headplate was affixed using dental cement. Postoperative care included the administration of carprofen (5 mg/kg, subcutaneously) daily for three days, and mice were monitored closely for signs of distress or weight loss. Post-surgery, animals were provided with a heat pad for 0.5–3 hours and offered softened chow to facilitate recovery.

For behavioral experiments, all mice were between 3 to 12 months old at the time of testing and were randomly assigned to experimental groups. Mice performing head-fixed treadmill-based dead reckoning tasks were housed individually post-surgery to prevent damage to the implanted headplate. Those participating in operant conditioning tasks, such as the effort-based lever press for reward-seeking behavior, were subjected to a gradual water restriction regimen to achieve a body weight reduction to 80–85% of their free-drinking weight. During the restriction period, they were habituated to the Skinner box for five minutes daily before formal training began.

Animals were systematically assigned to different experimental conditions to ensure balanced distribution across treatment groups. For tasks requiring drug or saline injections, the same mice were subjected to both pre-treatment baseline testing and post-treatment assessments, allowing for within-subject comparisons. To minimize stress and its potential confounding effects, all mice underwent gradual acclimatization to handling, head fixation, and experimental apparatuses prior to data collection.

## METHOD DETAILS

### Intramuscular and Intraperitoneal Injection

Intramuscular injections were performed under two different conditions: (1) under gaseous anesthesia and (2) in awake animals using physical restraint. Initial experiments were conducted under anesthesia using 3% isoflurane in 1.8 L/min pure oxygen for induction, followed by 1.5% isoflurane for maintenance. However, behavioral observations indicated that anesthesia significantly affected locomotion and cognitive performance for 10–30 minutes post-injection. To mitigate these effects, awake injections were introduced, wherein mice were physically restrained inside a multi-holed, funnel-shaped plastic bag. Each limb was carefully pulled through designated openings to immobilize the animal while minimizing stress.

A 100 μL Hamilton syringe fitted with a 30G needle was used to inject CTZ (coelenterazine, NanoLight, cat. no. 3031) dissolved in normal saline with 20 mM HEPES at a final concentration of 1 mg/mL. The injection site was determined based on preliminary observations of muscle activation and motor responses. Early experiments targeted the forearm muscles and gastrocnemius (distal limb muscles), but later injections were shifted to the triceps brachii, quadriceps femoris, and hamstring muscles (proximal limb muscles). This transition was based on the hypothesis that proximal muscle activation would provide stronger mechanical feedback relevant to self-motion perception. Injection volume was typically 30–50 μL per site, depending on the muscle size, and post-injection monitoring ensured that mice did not exhibit excessive struggling or distress.

### Headplate Implantation

For head-fixed treadmill experiments, a custom-designed stainless steel headplate was surgically affixed to the skull to facilitate stable fixation. Prior to surgery, the preoperative body weight was recorded to monitor postoperative recovery. Mice were anesthetized using 3% isoflurane in pure oxygen for induction, followed by 1.5% isoflurane at 1.2 L/min for maintenance. Once the respiratory rate slowed to approximately one breath per second, mice were secured in a stereotaxic frame with ear bars to ensure precise alignment during the procedure.

The surgical area was disinfected by alternating applications of 70% ethanol and povidone-iodine, and 50–100 μL of 2% lidocaine was injected subcutaneously at the incision site for local anesthesia. A midline incision was made to expose the skull, and all soft tissue was removed to ensure a clean surface for bonding. The headplate was positioned using a stereotaxic arm to ensure horizontal alignment and was affixed using dental cement (Metabond, Parkell). The cement was allowed to harden for 20 minutes before tapering isoflurane levels and monitoring the animal for recovery.

Postoperative care included administering carprofen (5 mg/kg, subcutaneously) daily for three consecutive days, and mice were monitored for signs of distress or weight loss. To facilitate recovery, mice were provided with heat pads for 0.5–3 hours and given softened chow soaked in water. Mice were housed individually after surgery to prevent damage to the headplate by cage mates.

### Dorsal Root Ganglia Isolation and Histological Sectioning

For immunohistochemistry and histological analysis, mice were deeply anesthetized with 3% isoflurane in 1.8 L/min pure oxygen and transcardially perfused with cold 1× phosphate-buffered saline (PBS), followed by 4% paraformaldehyde (PFA) in PBS. The spinal column was carefully dissected, and the dorsal root ganglia (DRG) were isolated according to the protocol described by Sleigh et al. (2016, 2020). Briefly, after removing the vertebrae, DRGs from cervical, thoracic, and lumbar regions were extracted using fine forceps and placed in 4% PFA at room temperature for 3 hours for fixation.

Following fixation, DRGs were transferred to 30% sucrose in PBS at 4°C overnight or until the tissue no longer floated, indicating complete cryoprotection. The DRGs were then embedded in Cryo-Gel embedding medium (Leica Surgipath) and frozen at −20°C. Cryosections were cut into 12 μm slices using a cryostat (Leica CM1950) and mounted onto SuperFrost Plus glass slides. Sections were allowed to dry overnight before immunostaining.

### Immunohistochemistry

Before staining, slides were washed in PBS to remove residual embedding medium. Sections were post-fixed in 4% PFA for 10 minutes, followed by three washes in PBS. Tissue permeabilization was performed using 0.2% Triton X-100 in PBS for 5 minutes, followed by three additional PBS washes. To block non-specific binding, sections were incubated in Animal-Free Blocking Solution (Vector Labs, USA) for 1 hour at room temperature. Primary antibody incubation was carried out overnight at 4°C in blocking solution. The following primary antibodies were used:

Anti-PV (mouse monoclonal, 1:1000, Swant, RRID: AB_10000343)

Anti-GFP (rabbit polyclonal, 1:1000, Abcam, ab290)

The next day, slides were washed three times with PBS before incubation with secondary antibodies for 3 hours at room temperature:

AF488 Anti-Rabbit IgG H&L (goat, 1:1000, Abcam, ab150077)

DL405 Anti-Mouse IgG (goat, 1:1000, Invitrogen, 35501BID)

Following secondary incubation, slides were washed three times in PBS and mounted using FluoroShield mounting medium with DAPI counterstaining.

### Microscopy and Image Analysis

Fluorescent images were acquired using a confocal microscope (Leica SP8) with 405 nm, 488 nm, and 561 nm laser lines for detecting DAPI, GFP, and PV signals, respectively. Image acquisition parameters were maintained across all samples to allow quantitative comparisons. For analysis, PV+ and LMO3+ cells in DRG slices were manually counted using Fiji (ImageJ).

### *In-Vivo* Bioluminescence Imaging

In vivo bioluminescence imaging was performed using an IVIS Spectrum 2 In Vivo Imaging System (Revvity) to assess the functionality of Luminopsin (LMO3). Before imaging, PV;LMO3 B6 transgenic mice and wild-type (WT) controls were anesthetized using 1.5–2% isoflurane in oxygen. The hair over the hindlimb muscles was removed with a depilatory cream to minimize signal scattering. Throughout the procedure, the body temperature of the mice was maintained at 37°C. A baseline image was acquired before the substrate injection. Subsequently, coelenterazine (CTZ; Nanolight #3031), dissolved in a HEPES-buffered saline solution, was administered via intramuscular (i.m.) injection into the proximal limb muscles at a dose of 8.2 mg/kg. Immediately after injection, mice were placed back into the imaging chamber. Dynamic bioluminescence images were acquired at 34.5-second intervals for a total of 40 shots, each with a 30-second exposure time.

### Freely Moving Open Field Test

To assess spontaneous locomotor activity, each mouse was placed in an open field arena consisting of four adjacent 40 cm × 40 cm acrylic boxes with white flooring and black walls. The testing room was illuminated by a ceiling-mounted fluorescent tube, providing an illumination of approximately 1000 lux at the center of each box. Before the experiment, mice were habituated to the testing room for 30 minutes to minimize stress-induced behavioral alterations. Following habituation, each mouse received an intramuscular injection of CTZ and was then immediately placed at the center of an open field arena. Mice were allowed to explore freely for 30 minutes, during which their movements were recorded using an overhead camera. To minimize external influences, the experimenter left the testing area during recording. After the session, the mouse was returned to its home cage, and the arena was thoroughly cleaned using 75% ethanol followed by a 500 ppm hypochlorous acid solution to eliminate residual odors. A waiting period of at least five minutes was enforced before introducing the next mouse to ensure the dissipation of any remaining cleaning agents.

### Rotarod Test

The rotarod test was used to evaluate motor coordination, balance, and endurance. The apparatus consisted of a horizontal rod that rotated at variable speeds. Before formal testing, mice were acclimated to the testing environment for 30 minutes to reduce stress. During the training phase, each mouse was placed on the rod, which rotated at a constant speed of 4 rpm for 60 seconds per trial. Three training trials were conducted per session, with a 10-minute inter-trial interval. Training sessions were repeated until the mice demonstrated stable performance. For the testing phase, mice were anesthetized using 3% isoflurane in 1.8 L/min oxygen, followed by intramuscular injection of saline (control) or CTZ (experimental group). After four minutes of recovery, the mouse was placed on the rotarod, which accelerated linearly from 0 to 40 rpm over a 300-second period. The time to failure was recorded, defined as the time when the mouse either fell off the rod or clung to it without actively walking. The test was conducted once per day over three consecutive days, with a two-day interval between each session to prevent fatigue effects.

### Effort-Based Lever-Press for Reward Seeking

To investigate motivation-based effort exertion, mice were trained to perform lever pressing for a sucrose reward in a custom-built operant chamber (10 cm × 10 cm × 10 cm, opaque acrylic walls with a transparent front panel). The setup consisted of a lever positioned 3 cm above the floor, a lick port, and an automated control system operated via an Arduino board. Mice were placed under water restriction (1 mL/day for at least three days) to reduce their body weight to 80–85% of their free-drinking weight before training. Magazine training was conducted to establish an association between lever pressing and sucrose reward (5 μL of 20% sucrose water per reward). Initially, mice received manually delivered rewards whenever they interacted with the lever (e.g., nose poking, touching, biting). Once they learned this association, they progressed to Fixed Ratio (FR) training, where pressing the lever a set number of times resulted in a reward. Mice first underwent FR1 training (one press per reward) and progressed to FR5 training (five presses per reward) once they achieved 100 rewards within 10 minutes for three consecutive days. The testing phase followed a Progressive Ratio (PR) schedule, where the effort required for each subsequent reward increased according to the equation:

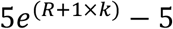

where R represents the number of rewards obtained, and k was set at 0.2 to balance difficulty. The session ended if the mouse either stopped pressing for 10 minutes, reached the daily water limit (1 mL), or exceeded one hour of testing. The breakpoint, defined as the highest ratio completed, was recorded daily. Prior to testing, mice were anesthetized with 3% isoflurane in 1.8 L/min oxygen before receiving an intramuscular injection of saline or CTZ. Immediately post-injection, the mouse was placed in the operant chamber to initiate the task. Lever pressing events were recorded using custom Arduino scripts and stored as CSV files for later analysis.

### Balance Beam Test

The balance beam apparatus consisted of a 1-meter-long flat beam of varying widths (24 mm, 12 mm, or 6 mm) positioned 70 cm above the floor with a safety net underneath to prevent injuries in case of falls. The beam connected a white start platform (15 cm × 10 cm) and a black home box (20 cm × 20 cm × 20 cm). To motivate mice to traverse the beam, a 30 W LED light was positioned above the start platform, creating a mild aversive stimulus, while the home box contained nesting material and food as an incentive. During the training phase, mice were trained to cross the 12-mm or 6-mm beam five times per session over two consecutive days. In the first trial, the mouse was allowed to rest for one minute upon reaching the home box. For subsequent trials, it was returned to the start platform after a 15-second interval. For the testing phase, mice received an intramuscular injection of either saline or CTZ under physical restraint before being placed at the start platform. Each mouse traversed both the 12-mm and 6-mm beams three times, with a 15-second interval between trials. Foot faults, defined as instances where the hindpaw completely slipped below the horizontal plane of the beam, were manually recorded using BORIS event-logging software.

### Grid Walking Test

Mice were placed on a 40 cm × 40 cm wire mesh grid, with each grid cell measuring 2 cm × 2 cm. A mirror (40 cm × 40 cm) was positioned beneath the grid at a 45-degree angle to allow simultaneous video capture of side and bottom views. The test was conducted in complete darkness to remove visual cues. All whiskers were trimmed 1–3 days before testing to eliminate tactile input variations. Following an intramuscular injection of saline or CTZ, mice were placed at the center of the grid, and video recordings were acquired at 120 FPS (1920 × 1080 pixels resolution) for 30 minutes. Bottom-view videos facilitated unambiguous identification of foot faults, while side-view recordings provided additional behavioral context.

### Head-Restrained Dead Reckoning Task

In this task, mice were head-fixed on a passive treadmill (Janelia 2017-049) and trained to associate distance-based cues with reward acquisition. Each trial began with a 7 kHz tone, signaling the start. The mouse ran from a virtual position of 0–5 cm toward a reward zone (150–170 cm), where a lick triggered a sucrose reward. If the mouse failed to lick within the reward zone, an inter-trial interval (ITI) of 0.5 seconds was enforced before resetting the virtual position. Training was divided into three phases: (1) random rewards, (2) fixed rewards at the reward zone, and (3) operant reward-triggering by licking. The test phase spanned four days, including a sham restraint session, a CTZ session, and a saline session, with one rest day in between. Running speed was recorded using a linear encoder, and licking behavior was logged with a capacitive touch sensor, with both signals stored in CSV files via Bonsai (C# scripts).

## QUANTIFICATION AND STATISTICAL ANALYSIS

### Open Field Test

Tracking and behavioral classification were performed under our CASTLE pipeline^57^, a training-free, region-of-interest(ROI)-informed hierarchical clustering algorithm and user package co-developed between Yu-Wei Wu Lab (Academia Sinica) and our group. The mouse was tracked by temporally aware video segmentation: the ROI was segmented with the Segment Anything Model (SAM) from Meta and propagated across all frames with DeAOT (*Decoupled Associating Objects with Transformers*) in CASTLE, and the body centroid was taken as the centroid of the tracked ROI mask. Behavior was then classified as “class” frame-by-frame from self-supervised DINOv3 features, which were embedded with hierarchically *human-in-the-loop* UMAP (the “Behavioral Microscope”) and then clustered into a 10-class ethogram, selected following careful cross-validations made between clustering robustness and human (expert) intuition at the level of raw ROI images (Walking, Running, Turn Left, Turn Right, Supported Rearing, Unsupported Rearing, Grooming, Sniffing, Immobility, and an unclassified “Others‒; class). Here, a “behavioral class” is a human-interpretable set of movie frames which has a longer dynamics in time than relatively short motor “motifs”.

### Centroid metrics

All single-window analyses used the 5–30 min segment; the first 5 min (dominated by a transient with the interferences from placement/handling) were excluded, and only analyzed separately in the time-course supplement (**Fig. S1**). The body-centroid trajectory was Gaussian-smoothed (σ ≈ 0.12 s) and downsampled to ∼30 fps. For each animal we computed the fraction of time moving (instantaneous speed > 4 cm/s, set above the ∼3.3 cm/s residual-jitter floor measured from Immobility-classified frames), total distance travelled, mean speed while moving, mean absolute acceleration while moving, and the fraction of time spent in the central zone of the arena (central 20 × 20 cm: |x − 20| < 10 and |y − 20| < 10 cm). From the ethogram, we computed the fraction of time for each behavior class and, per class, the median duration of behavior bouts (per animal). Spatial-occupancy maps were generated by binning the arena space into 1-cm² cells, normalizing each animal’s map to sum to 1, and averaging within each group (expressed in a log color scale).

### Velocity and acceleration vector fields

From the centroid trajectory (5–30 min) we computed per-step displacement vectors at a step of 0.2 s (∼5 Hz): velocity vectors and acceleration vectors. Vectors were pooled within each group and displayed as polar histograms (angle, movement direction; radius, magnitude, capped at the 99.5th percentile for display purposes; per-panel-normalized count, log color). Groups were compared at the animal level: each animal’s joint angle × magnitude polar histogram and spatial occupancy map were compared using an energy-distance statistic, and the significance was assessed by permuting animal group labels (1,000 resamples); the velocity and acceleration tests were FDR (False Discovery Rate)-corrected together.

### Statistics

Group comparisons for all single time window (5-30 min) metrics used *estimation statistics* (DABEST): the effect size is the mean difference (PV-Cre;LMO3 ^f/f^ − Control), reported with a 95% bias-corrected and accelerated (BCa) bootstrap confidence interval and a two-sided unpaired permutation p value (5,000 resamples). Multiple comparisons were controlled with the Benjamini–Hochberg False-Discovery-Rate procedure applied separately within three families: locomotor kinematics, class occupancy, and bout duration; statistical significance was taken at q < 0.05.

### Temporal dynamics (the time-course analysis supplement, Fig. S1)

To assess time dependence of the presumed luminopsin activation effects (e.g. CTZ washout and opsin kinetics), the full 0–30 min session was divided into six 5-min bins, and the same kinematic variables and class-occupancy metrics were computed per bin. Each metric was analyzed with an Aligned-Rank-Transform (ART) two-way repeated-measures ANOVA (factors: time × genotype; animal as the repeated measure)—the same procedure used for the Grid Walking analysis (see below), here implemented in Python (per-effect alignment, ranking, and then a mixed-design ANOVA on the ranks; only complete-bin animals were included for analyses). The genotype × time interaction was the primary effect of interest; the genotype and time main effect were reported in the source data. p values were FDR-corrected within the kinematics and behavior-class families separately.

### Rotarod Test

The time to failure (defined as the duration before the mouse either fell off or clung to the rod without actively walking) was manually recorded during testing. Data were analyzed using a two-way repeated measures ANOVA (RM ANOVA) in GraphPad Prism, with post-hoc multiple comparisons performed when significant interactions were found.

### Effort-Based Lever Press Test

The total number of lever presses and the breakpoint (the highest effort requirement completed before session termination) were extracted from Arduino-generated CSV files using a Python script. Statistical analyses were conducted in GraphPad Prism, with comparisons between groups performed using a two-tailed Mann–Whitney U test.

### Balance Beam Test

Behavioral annotation was conducted manually on a frame-by-frame basis using BORIS (Friard & Gamba, 2016) to record foot fault events. Data were plotted using a Python script, and statistical significance was assessed using an unpaired permutation test (5,000 iterations, DABEST) to compare foot fault frequencies across conditions.

### Grid Walking Test

Bottom-view video recordings were used to measure locomotion patterns and foot faults. DeepLabCut was used for tracking, with ResNet50 as the backbone model. Frames were selected for training using a k-means clustering approach, ensuring a well-balanced dataset. Keypoints were filtered using a 0.9 likelihood threshold, and travel distance normalization was applied to foot fault events. The temporal distribution of foot faults was analyzed using a Python script, and a non-parametric version of two-way RM ANOVA was conducted using Aligned Rank Transform (ART) (Wobbrock et al., 2011), followed by post-hoc permutation tests (5,000 iterations, DABEST).

### Head-Restrained Dead Reckoning Task

To quantify behavioral differences in spatial inference under LMO3 perturbation, three primary metrics were used:

Δ COM of lick positions from the reward zone (RZ) floor (150 cm), reflecting the difference between the animal’s inferred reward location and the actual reward location.

Δ Standard deviation of lick positions within a trial, indicating (empirical) certainty of spatial inference at a single-trial timescale.

Δ Standard deviation of COM of lick positions across trials, assessing (empirical) certainty across a 20-trial session.

For each mouse, trials 11–30 (Pre-Tx) served as baseline, and trials 41–60 (Post-Tx) were used for comparison. The first 10 trials following the treatment were excluded from analysis to account for potential transient effects.

Behavioral data were extracted from Arduino-logged CSV files and analyzed using custom Python scripts. Group comparisons were performed using an unpaired permutation test (5,000 iterations, DABEST), and within-subject effects were assessed using a paired permutation test.

### Analysis for Immunohistochemistry on The Dorsal Root Ganglia

For histological validation of LMO3 expression, PV+ and LMO3+ cells in DRG slices were manually counted using Fiji (ImageJ). Fluorescence intensity was quantified using automated thresholding methods, and co-localization analysis was conducted to assess the percentage of PV+ neurons expressing LMO3.

### *In-Vivo* Bioluminescence (IVIS) Imaging

Image analysis was conducted using Living Image® software. A region of interest (ROI) was manually drawn over the injection site on each limb. The bioluminescent signal was quantified and expressed as average radiance (photons/sec/cm²/sr). The data were background-subtracted using a control ROI from a non-signal area.

### Overall Statistical Framework and Visualization

Data visualization was performed using GraphPad Prism and Python (Matplotlib, Seaborn, DABEST). Most group comparisons used *estimation statistics*, an effect-size informed statistical estimation and visualization—the mean difference with a 95% bias-corrected and accelerated bootstrap (BCa) confidence interval, and an unpaired or paired permutation test (5,000 resamples). When many metrics were compared, Benjamini–Hochberg (BH) FDR correction across measure families was applied. Time-binned data (Open-Field and Grid-Walking time course) were analyzed with a non-parametric two-way repeated-measures ANOVA via Aligned Rank Transform (ART). Two-dimensional velocity and acceleration vector distributions were compared at the animal level with an energy-distance permutation test (1,000 permutations of group labels, FDR-corrected together). Where a simple two-group non-parametric test was used (e.g., Lever-Press metrics, Balance Beam foot faults), a two-tailed Mann–Whitney U or paired Wilcoxon test was applied. Statistical significance was set at p (or FDR q) < 0.05.

All statistical analyses were implemented in Python (SciPy, NumPy, Statsmodels, DABEST) and GraphPad Prism.

## ADDITIONAL RESOURCES

No additional resources beyond those listed in the Resource Availability and Key Resources Table sections were generated or utilized in this study.

However, to facilitate transparency and reproducibility, all custom-written code used for behavioral task control, data preprocessing, statistical analysis, and figure generation is publicly available. The code repository includes well-documented Python and C# scripts, along with a README file detailing repository structure, dependency installation, and step-by-step instructions for running the scripts.

The codebase can be accessed at:

GitHub Team repository: https://github.com/Ching-Lung-Hsu-Lab

For additional technical details or troubleshooting regarding the code, please contact the lead author at the email provided in the Resource Availability section.

## Supplemental Methods Details

In the lightweight VR system designed in house, the mouse was head-restrained over a treadmill. Behavioral outputs, such as licking and locomotion, were detected / recorded by a capacitive sensor and a linear treadmill encoder (implemented with a Teensy microchip) using an Arduino microcontroller. The signals were then relayed to *Bonsai*, a visual reactive programming framework. *Bonsai* enables parallel processing of heterogeneous data streams and allows for custom logic scripting. Using C# scripts in *Bonsai*, we calculated the animal’s position in the virtual space and its task state in real time (see below for the integration of *DeepLabCut-Live!*). These computations dynamically updated the VR environment custom-implemented in a game engine (*Unity*, which has been made prior to this project) with Arduino to provide feedback, including sucrose solution as rewards, to the animal.

The VR environment was displayed on screens positioned in front of the mouse (in some of our designs, through audio speakers), while cameras were set up to capture the animal’s behavior and facial movements in real time. In certain applications (which we did not apply to this project), behavioral videos were processed to estimate the animal’s pose using deep neural networks in *DeepLabCut-Live! (DLC-Live!)*. Critically, *DLC-Live!* had very short reaction time delays as we empirically measured (around tens of ms), and talked to *Unity* using a fast (and lossy, yet negligible) communication protocol (OSC). These pose estimations also enabled motion segmentation for further analyses. This closed-loop VR system continuously recorded behavioral data, updated the virtual environment, and delivered sensory feedback through screens, speakers or another device whenever applicable. For our particular implementations employed for this paper, we adopted only the “path-integration” part (without closed-loop video and audio feedback, except for the starting auditory cue). Additionally, the VR suite has been designed to integrate neural activity recording techniques, such as Ca²⁺ imaging, and extracellular and patch-clamp electrophysiological recording (see Wang et al., 2025). Hardware synchronization was achieved through Arduino triggers, ensuring alignment between behavioral, neural, and task-related data.

The task logic was represented in the visual-programming environment of *Bonsai* as a workflow, where individual colored circles corresponded to distinct C# scripts. In this *Bonsai* workflow, behavioral data were first read from Arduino using the *SerialReadLine* function. The data were then processed in a custom *GameRule* script to evaluate the game logic, which determined subsequent actions, such as playing a corresponding auditory cue (**Fig. 7C**).

## ACKNOWLEDGEMENTS

We thank Jen-Hau Yang (University of Connecticut) for his dedicated discussion and technical suggestions regarding the effort-based Lever-Press task. We also thank Yu-Shun Liu and Yu-Wei Wu (Academia Sinica) for their development of the CASTLE foundational-model-based classification, and guiding us for co-development regarding benchmarking, visualization and user-interface. We additionally appreciate Xian-Bin Huang, Yi Liu and Matthew Cheng (鄭丞言) for the early in-house development of the VR behavioral system and the closed-loop mechanism based on real-time computer vision. The funding was under generous support of the Institute of Biomedical Sciences (IBMS) and Career Development Award (CDA), as well as the I-AI-A (Innovative AI Applications in Humanities and Scientific Research Award) of Academia Sinica.

